# The Bric-à-Brac transcription factors are necessary for formation of functional germline stem cell niches through control of *dpp* expression in the *Drosophila melanogaster* ovary

**DOI:** 10.1101/689323

**Authors:** Laurine Miscopein Saler, Mathieu Bartoletti, Virginie Hauser, Anne-Marie Pret, Laurent Theodore, Fabienne Chalvet, Sophie Netter

**Affiliations:** Institute for Integrative Biology of the Cell (UMR9198); Brown University; Institute for Integrative Biology of the Cell; Paris Saclay Institute of Neurosciences

## Abstract

Many studies have focused on the mechanisms of stem cell maintenance *via* their interaction with a particular niche or microenvironment in adult tissues, but how formation of a functional niche is initiated, including how stem cells within a niche are established, is less well understood. Adult *Drosophila melanogaster* ovary Germline Stem Cell (GSC) niches are comprised of somatic cells forming a stack called a Terminal Filament (TF) and underlying Cap Cells (CCs) and Escort Cells (ECs), which are in direct contact with GSCs. In the adult, the Engrailed (En) transcription factor is specifically expressed in niche cells where it directly controls expression of the *decapentaplegic* gene (*dpp*) encoding a member of the Bone Morphogenetic Protein (BMP) family of secreted signaling molecules, which are key factors for GSC maintenance. In late third instar larval ovaries, in response to BMP signaling from newly-formed niches, adjacent primordial germ cells become GSCs. The *bric-à-brac* paralogs (*bab1* and *bab2*) encode BTB/POZ-domain containing transcription factors, that are also expressed in developing GSCs niches where they are required for TF formation. Here, we demonstrate that Bab1 and Bab2 display redundant cell autonomous function for TF morphogenesis and we identify a new function for these genes in GSC establishment. Moreover, we show that Bab proteins control *dpp* expression in otherwise correctly specified CCs, independently of En and its paralog Invected (Inv). In fact, our results also indicate that *en/inv* function in larval stages are neither essential for TF formation, nor GSC establishment. Finally, when *bab2* was overexpressed in ovarian somatic cells outside of the niche, where *en/inv* were not expressed, ectopic BMP signaling activation was induced in adjacent germ cells of adult ovaries, which formed GSC-like tumors. Together, these results indicate that Bab transcription factors are positive regulators of BMP signaling for acquisition of GSC status.

## INTRODUCTION

A stem cell niche or microenvironment allows, first, the establishment of stem cells, and second, the maintenance of a balance between stem cell self-renewal and differentiation. Much more is known about stem cell maintenance than about initial stem cell establishment. The interactions between niche and stem cells need to be strictly controlled for maintaining homeostasis of adult tissues. In fact, a defect in stem cell homeostasis can be pathological in humans, producing for example unregulated cancer stem cells (Plaks et al., 2015; Prager et al., 2019; Zhao et al., 2018), and may also be an important part of normal and premature aging when stem cell populations lose their potential to self-renew (Ermolaeva et al., 2018). The discovery of pre-metastatic niches in cancer (Aguado et al., 2017; Kaplan et al., 2005) also makes the study of the properties of stem cell niches a key for gaining headway in cancer biology.

The *Drosophila melanogaster* adult ovary has proven to be an excellent model for understanding how interaction with adjacent somatic niche cells allows for maintenance of Germline Stem Cell (GSC) status (Greenspan et al., 2015; Xie and Spradling, 1998). Approximately 20 individual GSC niches, each associated with a small number (2-3) of GSCs, are present in the *Drosophila* adult ovary at the tip of structures called germaria (Fig 1A). Each GSC niche is composed of several types of somatic cells: a Terminal Filament (TF), which is a stack of about 8 flattened cells, approximately 5 Cap Cells (CCs) present at the base of the TF (Gilboa, 2015), one triangularly-shaped transition cell connecting the TF and CCs (Panchal et al., 2017) and posterior to CCs, the anterior Escort Cells (ECs) (Fig 1A) (Wang and Page-McCaw, 2018). Both CCs and anterior ECs, are in direct contact with GSCs. CCs are considered to be the key component of GSC niches, their number correlating closely with the number of GSCs (Song et al., 2007; Xie and Spradling, 2000). CCs anchor GSCs to the niche by DE-cadherin-mediated adhesion (Song et al., 2002) and produce two Bone Morphogenetic Proteins (BMPs), Decapentaplegic (Dpp) and Glass Bottom Boat (Gbb), acting as short-range secreted signals required for GSC maintenance (Liu et al., 2010, 2015; Luo et al., 2017; Song et al., 2004; Wang et al., 2008; Xie and Spradling, 1998, 2000). Indeed, Dpp/Gbb signal is transduced in GSCs, via the presence of a specific receptor composed of both Thickveins and Punt, leading to phosphorylation of the transcription factor Mad (pMad), translocation of pMad into the nucleus and transcriptional repression of germline differentiation genes such as *bag-of-marbles* (*bam*) (Chen and McKearin, 2003b; Song et al., 2004). It was shown that the homeobox transcription factor Engrailed (En) binds *dpp cis*-regulatory sequences *in vitro* and activates *dpp* transcription in CCs (Luo et al., 2017). Several other signaling pathways, such as the JAK/STAT (Lopez-Onieva et al., 2008; Wang et al., 2008) and Hedgehog (Liu et al., 2015; Lu et al., 2015; Rojas-Ríos et al., 2012) pathways, are active in niche cells and have also been implicated in regulation of *dpp* expression in these cells. In addition, the ectopic expression of *dpp* or *engrailed* and ectopic activation of JAK/STAT signaling in somatic cells of the germarium, as well as the loss-of-function of *bam* in germ cells, all lead to a germarial GSC tumorous phenotype further supporting the implication of these factors in GCS homeostasis (Eliazer et al., 2014; McKearin and Ohlstein, 1995; Song et al., 2004; Wang et al., 2008; Xie and Spradling, 1998).

**Fig 1.**
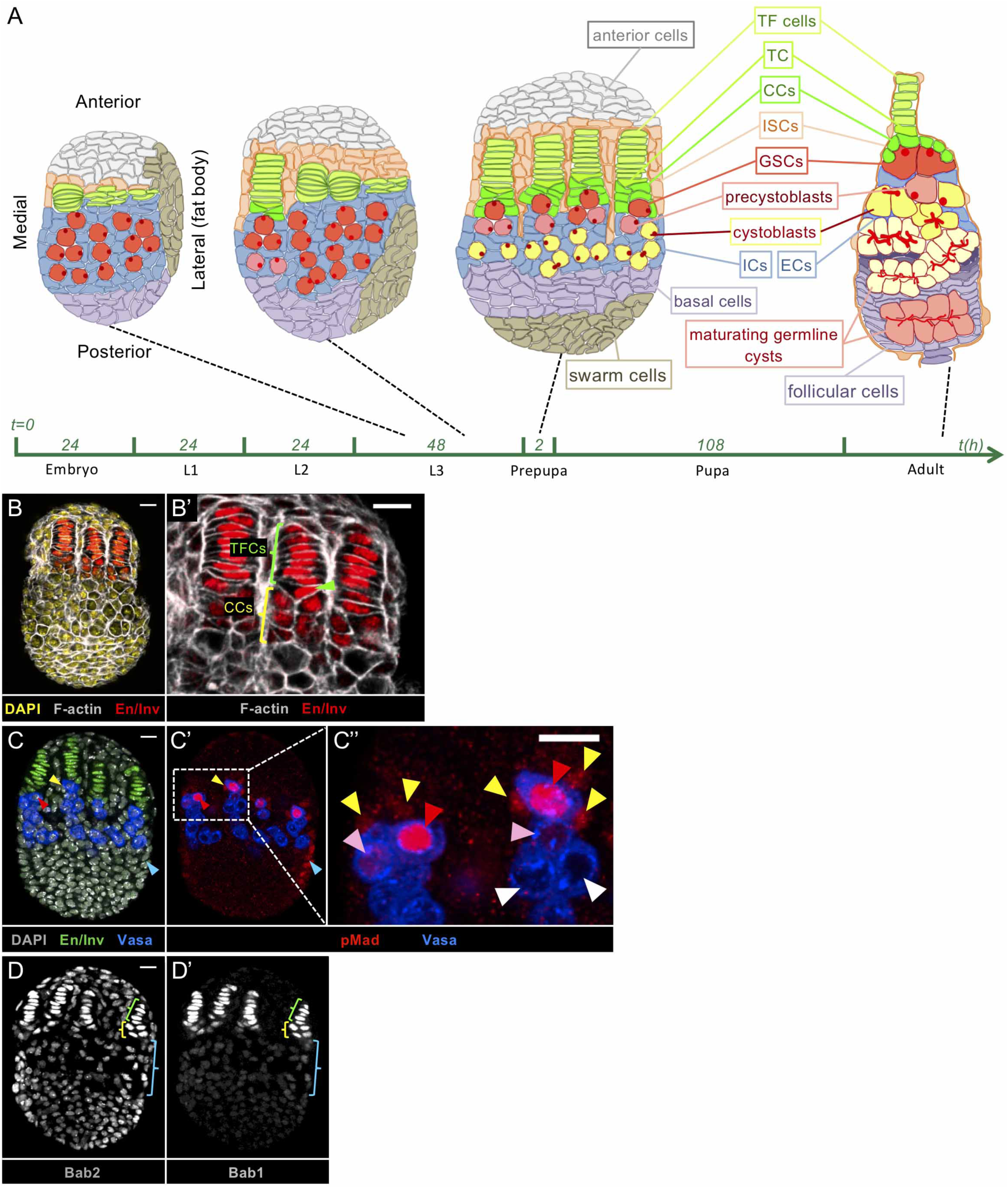
Specific cell types of the developing *Drosophila* ovary. (A) Schematic drawings of a developing ovary from mid L3 to prepupal stages and of an adult germarium, with all cell types indicated and represented by a different color. Anterior is to the top and for the developing ovary, medial is to the left. TF cells: Terminal Filament cells. TC: Transition Cell. CCs: Cap cells. ISCs: Inner Sheath Cells. GSCs: Germline Stem Cells. ICs: Intermingled Cells. ECs: Escort cells. The round spherical structure in germ cells during larval and prepupal stages represent the spectrosome and the red branched structures in germ cell cysts in the germarium represent fusomes, which are derived from spectrosomes. Spectrosomes and fusomes are cytoplasmic, cytoskeletal structures, and are used as specific germ cell markers. (B-D’) Whole mount immunostaining of wild type prepupal ovaries (projections of adjacent confocal sections). Anterior is up, medial is to the left. Scale bar: 10 µm. (B’,C’’) Higher magnification of the niche region of the corresponding ovary. (B,B’) Engrailed/Invected (En/Inv) (red) mark the niche cell nuclei specifically and F-actin labeling (grey) marks all cell membranes. TF cells (green bracket) are characterized by a flat shape, with flat nuclei accumulating high levels of En/Inv, while CCs (yellow bracket) are cuboidal, with weaker En/Inv expression level. The triangular TC (green arrowhead) is located between the TF cells and the CCs. (C) En/Inv (green) marks the nuclei of niche cells, and Vasa (blue) marks the cytoplasm of Germ Cells (GCs). (C’,C”) In the most anterior GCs, in contact with niche cells, the BMP signaling pathway is activated as evidenced by accumulation of pMad (red) at high (red arrowheads) or low (pink arrowheads) levels. GCs accumulating pMad and in contact with CCs are considered as GSCs. CCs (yellow arrowheads) and other peripheral posterior cells (blue arrowhead) accumulating pMad (red) can be differentiated from GCs since they do not accumulate Vasa (blue). The most posterior GCs, engaged into differentiation, do not accumulate pMad (white arrowheads). (D) Bab2 accumulates in the nuclei of all somatic cells but at a higher level in niche TF cells and CCs (green and yellow brackets, respectively, compared to blue bracket). (D’) Bab1 accumulates in niche TF cells and CCs and is also reported here for the first time to be present at low levels in ICs at this stage (blue bracket).

Morphogenesis of GSC niches is a highly stereotyped process occurring in the third instar larval ovary. Morphogenesis begins with TF formation which involves flattening, sorting, intercalation and stacking of somatic TF cell precursors, initiating at the medial side of the ovary and progressing as a wave laterally (Fig 1A) (Godt and Laski, 1995; Sahut-Barnola et al., 1996). Individualization of each TF is accomplished by the migration of apical somatic cells (Inner Sheath Cells) between TFs (Fig 1A) (Cohen et al., 2002).The number of TFs that form in the larval ovary determines the number of GSC niches at the adult stage (Bartoletti et al., 2012; Green et al., 2011; Sarikaya et al., 2012). At the base of each newly-formed TF, the anterior-most Intermingled Cells (ICs-somatic cells intermingled with Primordial Germ Cells, PGCs) differentiate into CCs adopting a cuboidal shape clearly distinguishable from that of the flat TF cells (Fig 1A and Fig 1B and B’, yellow and green brackets, respectively) (Panchal et al., 2017; Zhu and Xie, 2003). At this stage, the transition cell is already distinguishable (Fig 1A and 1B’, green arrowhead) (Panchal et al., 2017). Posterior-most ICs will give rise to ECs in adult ovaries (Fig 1A) (Lai et al., 2017).

Very few genes have been reported to be implicated in TF formation, notably, the *bric-à-brac* locus paralogs (*bab1/bab2*) (Bartoletti et al., 2012; Couderc et al., 2002; Godt and Laski, 1995; Green and Extavour, 2012; Sahut-Barnola et al., 1996) and the *engrailed*/*invected* locus paralogs (*en/inv*) (Bolívar et al., 2006; Gustavson et al., 1996). The *bab1* and *bab2* genes encode proteins sharing evolutionarily conserved domains: a BTB (Broad-Complex, Tramtrack and Bric-à-brac)/POZ (POx virus and Zinc finger) domain involved in homodimeric and heterodimeric interactions (Chaharbakhshi and Jemc, 2016) and a Bab-Conserved Domain (BabCD) containing two known motifs, a Pipsqueak (Psq) domain and an AT-hook like motif both involved in DNA-protein interactions (Couderc et al., 2002). In the larval ovary, loss of function alleles of the *bab* locus lead to a dominant phenotype characterized by an excess of TFs, resulting in an excess of GSC niches in adults (Bartoletti et al., 2012), and by a recessive phenotype characterized by a defect in TF formation resulting in atrophied ovaries with few germ cells in adults and sterility (Godt and Laski, 1995; Sahut-Barnola et al., 1995). As for the *engrailed/invected* locus, induction of clones homozygous for a deletion encompassing both genes identified a cell autonomous function for these genes in the correct alignment of TF cells during TF formation (Bolívar et al., 2006). CC specification has been shown to require the combined action of Notch signaling and the large Maf transcription factor Traffic Jam (Tj) (Gancz et al., 2011; Hsu and Drummond-Barbosa, 2011; Panchal et al., 2017; Song et al., 2007; Yatsenko and Shcherbata, 2018).

The newly-formed niches can be considered functional at the prepupal stage when the underlying PGCs become GSCs, that is, when they begin to undergo asymmetric divisions producing two different types of daughter cells, a GSC that remains within the niche and a cystoblast that will give rise to a germline cyst (Zhu and Xie, 2003). Before niche formation, that is, before the mid third instar larval stage, all PGCs have been shown to exhibit BMP signaling activation (Gilboa and Lehmann, 2004; Kai and Spradling, 2004), and this is correlated with detection of Dpp in all somatic cells of the ovary (Sato et al., 2010). During the larval to pupal stages, Dpp signaling prevents PGC differentiation into germline cysts (Gilboa and Lehmann, 2004; Matsuoka et al., 2013; Zhu and Xie, 2003) and promotes PGC proliferation (Sato et al., 2010; Zhu and Xie, 2003). Upon niche formation, *dpp* expression (Matsuoka et al., 2013; Zhu and Xie, 2003) and Dpp protein accumulation (Sato et al., 2010), become unevenly distributed in the ovary by an as yet uncharacterized mechanism, with highest levels found in niche cells. Among GCs, only those in contact with the Dpp-secreting niche cells retain BMP signaling activation as evidenced by the presence of pMad (Fig 1C and 1C”, red and pink arrowheads) (Gancz et al., 2011) and become GSCs that retain the capacity to divide (Zhu and Xie, 2003). The more posterior PGCs, not in contact with niche cells, lose Dpp signaling activation (Fig 1C-1C”, white arrowheads) and proceed to differentiate into cystoblasts, whereupon they produce the first germline cysts (Matsuoka et al., 2013; Tseng et al., 2018; Zhu and Xie, 2003).

In this study, we addressed the role of *bab1* and *bab2* in formation of functional GSC niches, in particular, on establishment of the first GSCs in the larval ovary. In developing ovaries, Bab1 has been reported to be present only in niche cell and Bab2 in all somatic cells, however at higher levels in niche cells (Couderc et al., 2002). We also asked whether the two *bab* genes have distinct or redundant functions within developing niche cells for either niche formation or GSC establishment. For this, we used a null allele of *bab1* (*bab^A128^*) and Gal4-RNAi targeted knockdown of each or both of the two paralogs specifically in GSC niches during their formation in larvae. This approach has allowed us to show that *bab1* and *bab2* display redundant cell autonomous functions for TF formation. In contrast, cells exhibiting several CC characteristics were nonetheless present upon depletion of Bab proteins in niches. In addition, we have identified a new essential role for *bab* genes in prepupal niches for initial establishment of GSCs correlated with a role in *dpp* expression in CCs. The function of *bab* in CCs is unlikely to require that of the *en*/*inv* genes since their expression levels are not affected when Bab1 and Bab2 are depleted. Unexpectedly, we also found that En/Inv are not essential for activation of BMP signaling and GSC establishment in the larval ovary, contrasting with their essential functions in adult ovaries for BMP-mediated GSC maintenance. Finally, when *bab2* was overexpressed in somatic ovarian cells as of larval stages, GSC-like tumorous germaria were produced in the adult ovary. This was associated with ectopic BMP signaling activation occurring in GCs without the presence of En/Inv in the *bab2*-overexpressing somatic ovarian cells. Together, our results indicate that the *bab* genes as necessary for *dpp* expression in the newly-forming GSC niches and for establishment of the first GSCs.

## RESULTS

### *bab1* and *bab2* share redundant cell autonomous functions for TF morphogenesis

We addressed the effect of depletion of Bab1 and Bab2 proteins, either separately or together, specifically in two niche cell populations, TF cells and CCs, during niche formation at the third instar larval stage. First, we tested and validated two strategies based on the use of RNAi and/or mutant alleles to deplete each of these two proteins specifically in TFs and CCs. Bab2 is detected in all ovarian somatic cells at this stage, but at higher levels in TFs and CCs as previously reported (Couderc et al., 2002) (Fig 1D, green and yellow brackets, respectively). We used a *hedgehog-Gal4* (*hhG*) driver in order to express a *bab2*-specific RNAi construct (*UAS-bab2^IR^*) only in niche cells during their differentiation (Lai et al., 2017; Sarikaya and Extavour, 2015). In parallel, *hhG* was also used to drive a *UAS-GFP* reporter construct (*hhG>GFP*) to mark niche cells (thereafter named *hhG+* cells). In control *hhG>GFP* prepupal ovaries, *hhG* driver expression was found to be specific to niche TF cells and CCs, with much higher GFP accumulation in medial vs. lateral niches (Fig 2A and 2A’, yellow vs. blue dotted lines, respectively, and 2A”). Inside each niche, *hhG>GFP* expression levels were mosaic between TF cells and always low in CCs (Fig 2A’). Consistent with the expression profile of the *hhG* driver (Fig 2B’ and 2B”), in *hhG>UAS-bab2^IR^* ovaries, Bab2 depletion was observed in all medial TF cells, but not in all CCs (Fig 2B’, yellow dotted lines), and was not observed in lateral *hhG+* cells (Fig 2B’, blue dotted lines). Noteworthily, in niche cells knocked down for *bab2*, the level of Bab1 was not noticeably affected (Fig 2B’). Bab1 accumulation in the control *hhG>GFP* ovaries was detected at high levels in both TF cells and CCs as reported (Couderc et al., 2002) (Fig 1D’, green and yellow brackets, respectively). In addition, low level of Bab1 were also detected in ICs (Fig 1D’, blue bracket). For *bab1* depletion, we used *bab^A128^*, reported as a null mutant allele of *bab1* not affecting *bab2* expression (Couderc et al., 2002; Godt et al., 1993). Indeed, in prepupal ovaries from females homozygous for *bab^A128^*, *bab1* expression was undetectable, whereas the level of Bab2 did not appear to be different from that in the *hhG>GFP* control (Fig 2C and C’).

**Fig. 2.**
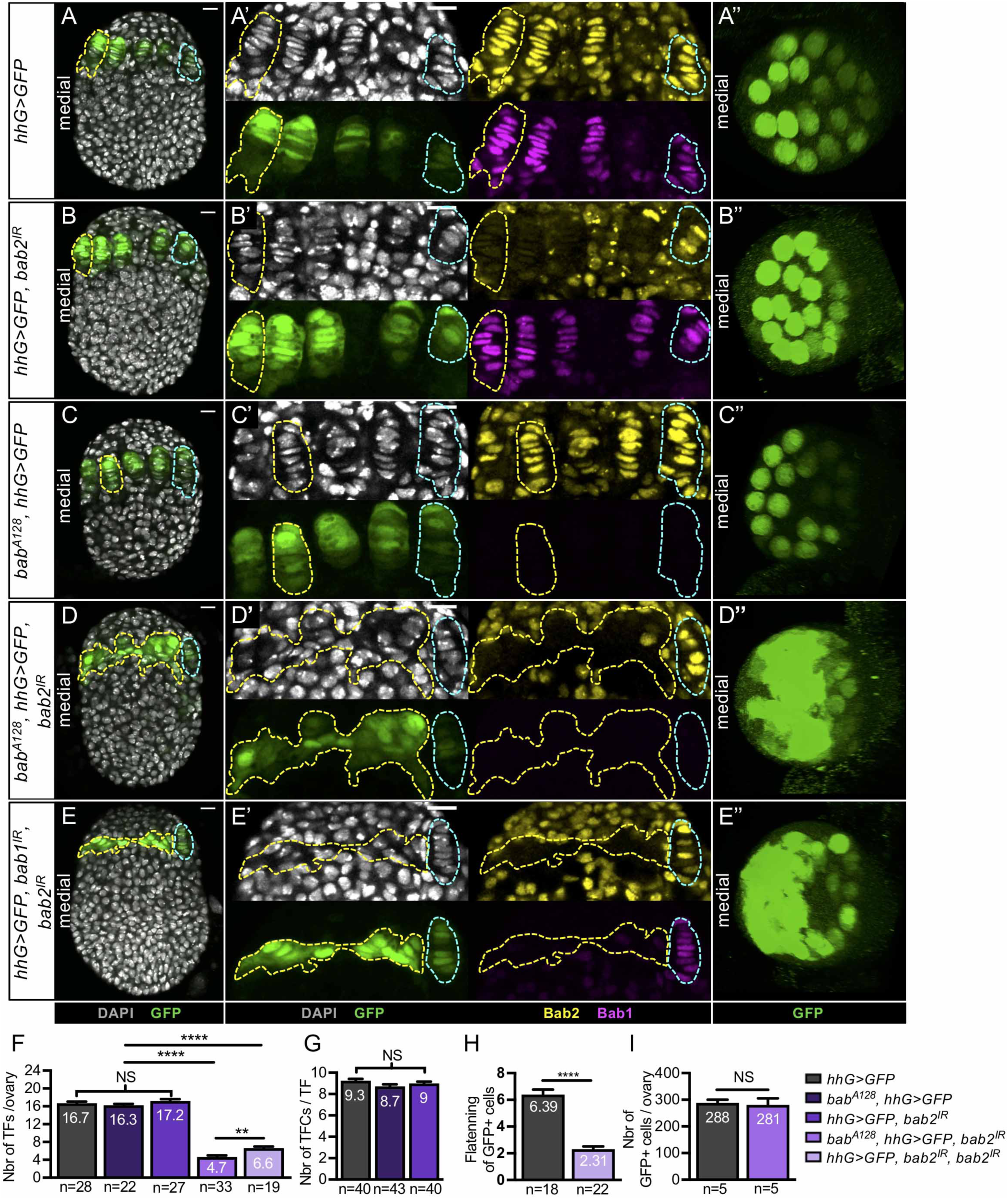
Depletion of Bab1 and Bab2 together in niches of larval ovaries impedes Terminal Filament formation. (A-E) Whole mount immunostaining of prepupal ovaries (projections of adjacent confocal sections) for detection of GFP (green), Bab1 (pink) and Bab2 (yellow). Nuclei are labeled with DAPI (grey). Anterior is up, medial is left. Scale bars: 10 µm. (A’,B’,C’,D’,E’) Higher magnification of the niche region of the corresponding ovaries. (A’’,B’’,C’’,D’’,E’’) Anterior view of ovaries using 3D reconstruction. Each GFP (green) circle corresponds to one Terminal Filament (TF) cross section. (A) Control ovary expressing a *UAS-GFP* construct under the control of a *hedgehog(hh)-Gal4 driver (hhG>GFP).* This driver is expressed as a decreasing medial (yellow dotted lines) to lateral (blue dotted lines) gradient in the TFs and CCs of niches as evidenced by the GFP (green) reporter. Moreover, in a single niche, the expression pattern of *hhG* is mosaic between TF cells and is always low in CCs at the base of each TF. Bab2 (yellow) accumulates in nuclei of all somatic ovarian cells, but at a higher level in TF cell and CC nuclei. Bab1 (magenta) accumulates at a high level in TF cell and CC nuclei. (B) Prepupal ovary expressing RNAi against *bab2 (hhG>GFP, bab2^IR^).* Consistent with the expression profile of the *hhG* driver, Bab2 depletion is obtained in all medial TF cells but not in all CCs at the base of these TFs (yellow dotted lines), and is not obtained in lateral *hhG+* cells (blue dotted lines). Bab1 (magenta) accumulation is not noticeably affected. (C) Prepupal ovary mutant for *bab1* (*bab^A128^, hhG>GFP).* Bab1 (magenta) is not detectable in medial or lateral niches, and the level of Bab2 (yellow) is not noticeably affected. For both genetic contexts (B,C), the morphology of the TFs and TF cells appear unaffected compared to the control (A). (D) Prepupal ovary expressing RNAi against *bab2* coupled with a homozygous *bab1* null mutation (*bab^A128^; hhG>GFP, bab2^IR^).* As in B, Bab2 (yellow) depletion is obtained in medial (yellow dotted line) but not lateral *hhG+* cells (blue dotted line) and Bab1 (magenta) is absent. (E) Prepupal ovary expressing RNAi against *bab1* and *bab2* (*hhG>GFP, bab1^IR^, bab2^IR^).* Consistent with B, Bab1 (magenta) and Bab2 (yellow) depletion is obtained in medial (yellow dotted line) but not lateral *hhG+* cells (blue dotted line). For both genetic contexts (D,E), in the medial part of the ovary (yellow dotted lines), the *hhG+* cells fail to flatten, stack and form TFs. (F-I) Graphs comparing different parameters related to TF formation, in control ovaries (*hhG>GFP*) and in the ovaries depleted of Bab1 and/or Bab2. (F) The number of TFs per ovary is not significantly different between the control and the ovaries depleted of Bab1 or Bab2. However, the ovaries depleted of both Bab1 and Bab2 have significantly fewer TFs per ovary than the control. (G) The number of TF cells (TFCs) per TF is not different between control ovaries and ovaries depleted of Bab1 or Bab2. (H) Control *hhG+* cells are significantly flatter than those knocked down for *bab1* and *bab2*, with flattening measured as the ratio of the width and height. (I) The total number of *hhG+* cells is comparable in control ovaries and ovaries depleted of Bab1 and Bab2 (*bab^A128^; hhG>GFP, bab2^IR^*). Values are presented as means +s.e.m. p-values are calculated using a one-way ANOVA test for F and G and a two-tailed t-test for H and I. n: sample size; NS: Not Significant (p>0.05); ****: (p<0.0001).

In either the *hhG>UAS-bab2^IR^* or the homozygous *bab^A128^* context, use of *hhG>GFP* to mark TFs indicated that the morphology of TFs and TF cells (Fig 2B and 2B’ and 2C and 2C’), the number of TFs per ovary (Fig 2F) and that of TF cells per TF (Fig 2G) were not different from that in the control. This was also the case when a *UAS-dicer2* construct was used to increase the effect of the *UAS-bab2^IR^* construct (Fig 3C-3C”). From these results, we concluded that loss of function of *bab1* alone or a strong depletion of Bab2 protein in TF cells did not affect TF morphogenesis.

**Fig 3.**
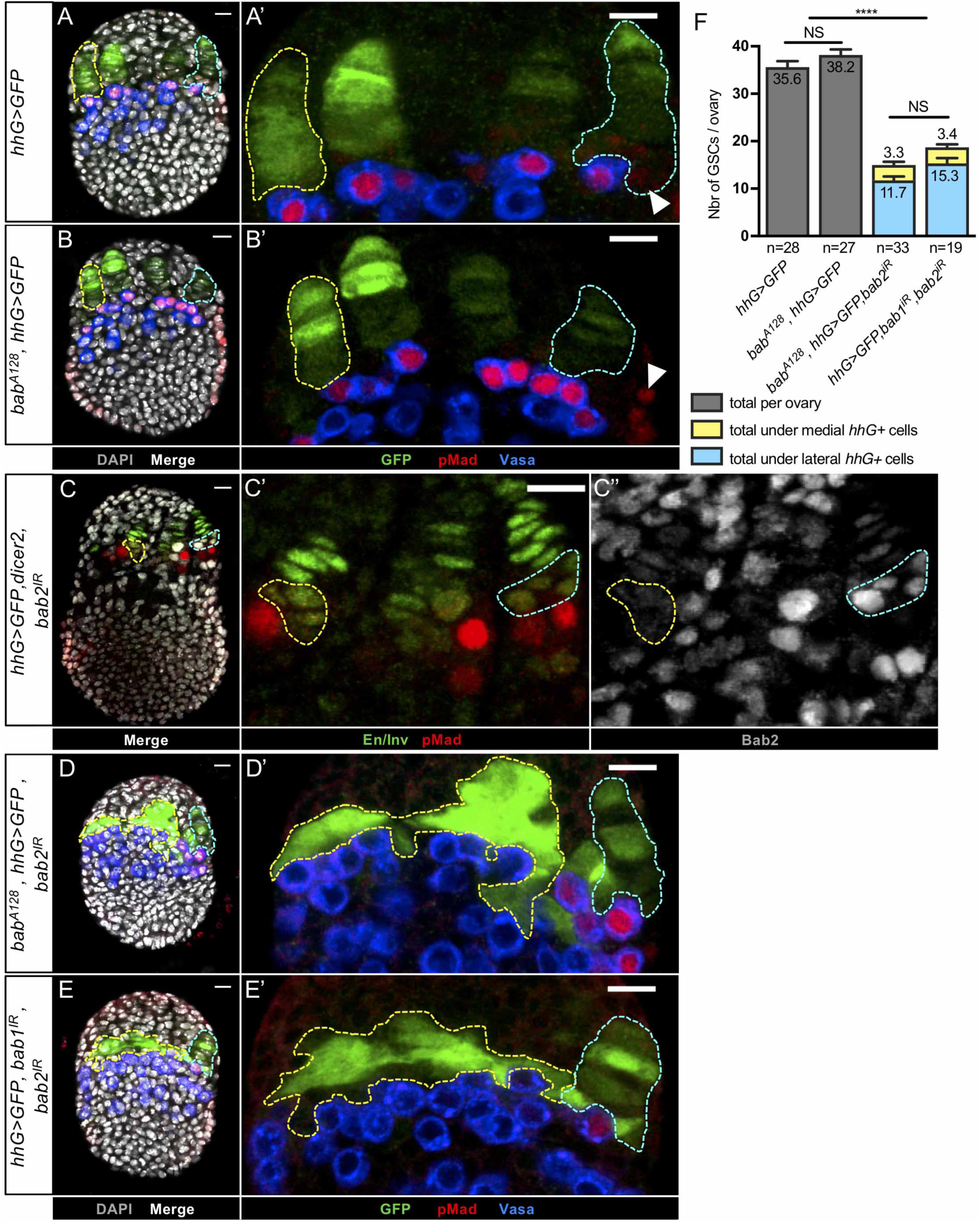
Bab1 and Bab2 depletion in developing niches impedes Germline Stem Cell establishment. (A-E) Whole mount immunostaining of prepupal ovaries (projections of adjacent confocal sections). Nuclei are labelled with DAPI (grey). Anterior is up, medial is left. Scale bars: 10 µm. The yellow dotted lines encircle the medial Terminal Filament (TF)/region, and the blue dotted lines, the lateral TF of each ovary. The transgenes and mutations present are indicated to the left of each row. (A’,B’,C’,C”,D’,E’) Higher magnification of the niche region (GFP, green) of the corresponding ovary. (A,A’) In a control ovary, pMad (red) immunostaining is detected in the anterior-most Germ Cells (GCs), marked with the presence of Vasa protein (blue). These Vasa+/pMad+ are in direct contact with the niches and are considered as Germline Stem Cells (GSCs). (B,B’) Ovary mutant for *bab1* (*bab^A128^; hhG>GFP)*. No significant difference was observed when compared to the control, regarding the presence of Vasa+/pMad+ GSCs in contact with the niches. (A’,B’) The CCs and some peripheral posterior cells (arrowheads) accumulate pMad (red), but can be differentiated from GCs since they do not accumulate Vasa (blue). (C-C”) Ovary expressing *UAS-nlsGFP, UAS-bab2^IR^* and *UAS-dicer2* under control of *hedgehog(hh)-Gal4 (hhG>GFP, dicer2, bab2^IR^).* In both the medial part of the ovary, where depletion of Bab2 protein (grey) is strong in CCs (C”, yellow dotted lines), and in lateral niches where it is not (C”, blue dotted lines), GSCs, marked by nuclear pMad staining (red), are associated with CCs. (D-E) Ovaries depleted of Bab1 and Bab2 using two genetic contexts*, bab^A128^; hhG>GFP, bab2^IR^* (D) and *hhG>GFP, bab2^IR^, bab2^IR^* (E). In the medial part of the ovary where we have shown that Bab protein depletion is strong, *hhG+* cells do not form TFs (yellow dotted lines), Vasa+ GCs (blue) are present, but almost none of them are GSCs since they are not pMad+. However, in the lateral region, where we have shown Bab protein depletion is weak, normal TFs are formed (blue dotted lines) and these are associated with Vasa+/pMad+ GSCs. (F) Graph comparing the overall number of GSCs per ovary in control ovaries and in ovaries depleted of Bab1 alone (*bab^A128^*) or both Bab1 and Bab2 using the two different genetic contexts. A significantly lower number of GSCs per ovary were obtained only when Bab1 and Bab2 were depleted together compared to the control, with no significant difference between the two mutant contexts. Moreover, the majority of GSCs in both mutant contexts were found in the lateral part of the ovary were TFs from (blue bar) and very few in the medial part (about 3 GSC per ovary, yellow bar). Values are presented as means +s.e.m. p-values are calculated using a one-way ANOVA test. n: sample size; NS: Not Significant (p>0.05); ****: (p<0.0001).

Next, *bab1* and *bab2* were knocked down together, in the *bab^A128^, hhG> GFP*, *bab2^I R^* or in the *hhG> GFP, bab1^IR^, bab2^IR^* contexts. In *bab^A128^, hhG> GFP, bab2^IR^* prepupal ovaries (Fig 2D), Bab1 protein was absent from all *hhG+* cells (Fig 2D’, yellow and blue dotted lines, respectively) and ICs as expected in *bab^A128^* mutants. In the same ovaries, Bab2 protein was efficiently depleted in medial but not lateral *hhG+* cells (Fig 2D’, yellow vs. blue dotted lines). In *hhG>GFP, bab1^IR^, bab2^IR^* ovaries (Fig 2E), *bab1* and *bab2* were only efficiently knocked down in medial *hhG+* cells, but not in lateral *hhG+* cells (Fig 2E’, yellow vs. blue dotted line, respectively). In both genetic contexts, the medial part of the ovary contained a large cluster of *hhG+* cells, which, unlike in the control, failed to form TFs (Fig 2D’ and 2E’, yellow dotted line). These results are consistent with the previously-described phenotypes of *bab* locus mutants (Godt and Laski, 1995). We quantified the number of TFs per ovary (Fig 2F), the flattening of the medial *hhG+* cells (Fig 2H) and found that these parameters were significantly lower than those in the control, while the overall number of *hhG+* cells per ovary was the same as that in the control (Fig 2I). In contrast, in the lateral-most part of the ovary, where *hhG* driver expression was lower and incomplete depletion of Bab proteins was obtained (Fig 2D-D” and 2E-E”), *hhG+* cells formed TFs that were not different from control lateral TFs (Fig 2D’ and 2E’ compared to 2A’, blue dotted lines). The higher penetrance of the abnormally low TF number per ovary phenotype using *bab^A128^* instead of *UAS-bab1^IR^* for the double *bab1/bab2* knockdown (Fig 2F) may be attributable to the more efficient depletion obtained for *Bab1* with *bab^A128^* than with *UAS-bab1^IR^* (Fig 2D’ compared to 2E’). Since reduction of both Bab1 and Bab2 in niche cells impaired TF formation significantly, while reduction of each one individually did not, we conclude that *bab1* and *bab2* share redundant, cell autonomous functions for TF morphogenesis.

### The functions of *bab1* and *bab2* are necessary in niche cells for GSC establishment in larval ovaries

As shown above, although *hhG+* cells depleted of Bab1 and Bab2 do not form morphologically normal niches, the normal number of *hhG+* cells are present as a mass of cells in the medial part of the ovary. We asked whether this un-organized *hhG+* cell mass was able to recruit the initial GSCs in the prepupal ovary. The same approaches as above were used to deplete Bab1 and Bab2 proteins from niche cells, either separately or together. To test for functional niche activity, BMP pathway activation in GCs was used as a characteristic of GSC identity at this stage. For this, accumulation of a direct downstream component of BMP pathway activation, phosphorylated Mad (pMad), was monitored in GCs marked by the presence of Vasa. In control prepupal ovaries, pMad was detected in anterior-most GCs adjacent to niches at either high or medium levels (Fig 1C’ and 1C”, red and pink arrowheads, respectively, and 3A and 3A’). In addition, pMad was also detected in two somatic cell populations, in particular, at a low level in CCs and at an intermediate level in peripheral posterior cells (Fig 1C’ and 1C”, yellow and blue arrowheads, respectively and Fig 3A’ and 3B’, white arrowheads, respectively). This is consistent with a previous report indicating BMP signaling activation in somatic cells using a different reporter (Zhu and Xie, 2003). In a *bab1* homozygous null mutant background (*bab^A128^, hhG>GFP*), pMad+ GSCs were present at the base of TFs (Fig 3B and 3B’) and in the same number as that in the control (Fig 3F). In *hhG>GFP, dicer2, bab2^IR^* ovaries (Fig 3C), very rare niches containing only CCs with undetectable levels of Bab2 were obtained (Fig 3C’ and C”, yellow dotted line). However, in these cases, GCs that were adjacent to Bab2-depleted CC groups (yellow dotted line) were always positive for pMad (Figure 3C’ and 3C’’). These results suggest that the depletion of either Bab1 or Bab2 individually in niche cells did not affect GSC establishment.

Upon strong depletion of both Bab1 and Bab2 in either *bab^A128^, hhG> GFP bab2^IR^* or *hhG> GFP, bab1^IR^, bab2^IR^* ovaries, Vasa+ GCs were found in close contact with the medial *hhG+* cells, but almost all GCs were devoid of pMad staining (Fig 3D and 3D’ and 3E and 3E’, below yellow dotted line). In contrast, in the lateral region where TFs were formed, pMad^+^ GCs (i.e. GSCs) were found at the base of each niche as observed in control ovaries (Fig 3D and 3D’, and Fig 3E and 3E’, bellow blue dotted lines). The overall number of pMad^+^ GCs per ovary was significantly reduced when compared to that in the control and, when present, these GSCs were associated with lateral TFs in almost all of cases (Fig 3F). Thus, in absence of both *bab1* and *bab2* functions, GCs were present and correctly positioned next to *hhG+* cells, but the BMP pathway was, for the most part, not activated in these cells, indicating that GSC status was compromised. We next asked whether these GCs may differentiate precociously. Germline cell differentiation can be monitored by the expression of a *GFP* transcriptional reporter for *bam* expression (*bamP-GFP*) (Chen and McKearin, 2003a). In control prepupal ovaries (*bamP-GFP, hhG>lacZ*), GCs (GFP+/Vasa+) that differente were only rarely detected among GCs contacting *hhG>UAS-lacZ (β-Gal+)* niche cells, whereas all GCs located one cell diameter away from these niche cells expressed this differentiation marker (Fig 4A and 4A’, Vasa+ cells under encircled TFs, and Fig 4C). In contrast, differentiating GCs (GFP+/Vasa+) were found significantly more frequently in direct contact with clusters of disorganized *hhG>UAS-lacZ (β-Gal+)* cells depleted of Bab1 and Bab2 in the medial zone of prepupal ovaries than in control ovaries (Fig 4B and 4B’, arrowhead, and Fig 4C). Abnormal differentiation of some GCs adjacent to the cluster of medial *hhG+* cells thus correlated with the abnormal absence of the GSC marker pMad in the same region. Taken together these results indicate that *bab* function is required in niches for acquisition of GSC status by PGCs as marked by sustained activation of the BMP pathway and absence of expression of the differentiation reporter *bamP-GFP*. Our results also indicate that Bab1 and Bab2 depletion in *hhG+* cells does not hinder GCs from remaining in close contact with *hhG+* cells, indicating that the absence of BMP pathway activation in these GCs cannot be attributed to an incapacity of these cells to contact niche cells. Together, our work strongly suggest that *bab1* and *bab2* share redundant functions for GSC establishment in the late larval ovary.

**Fig 4.**
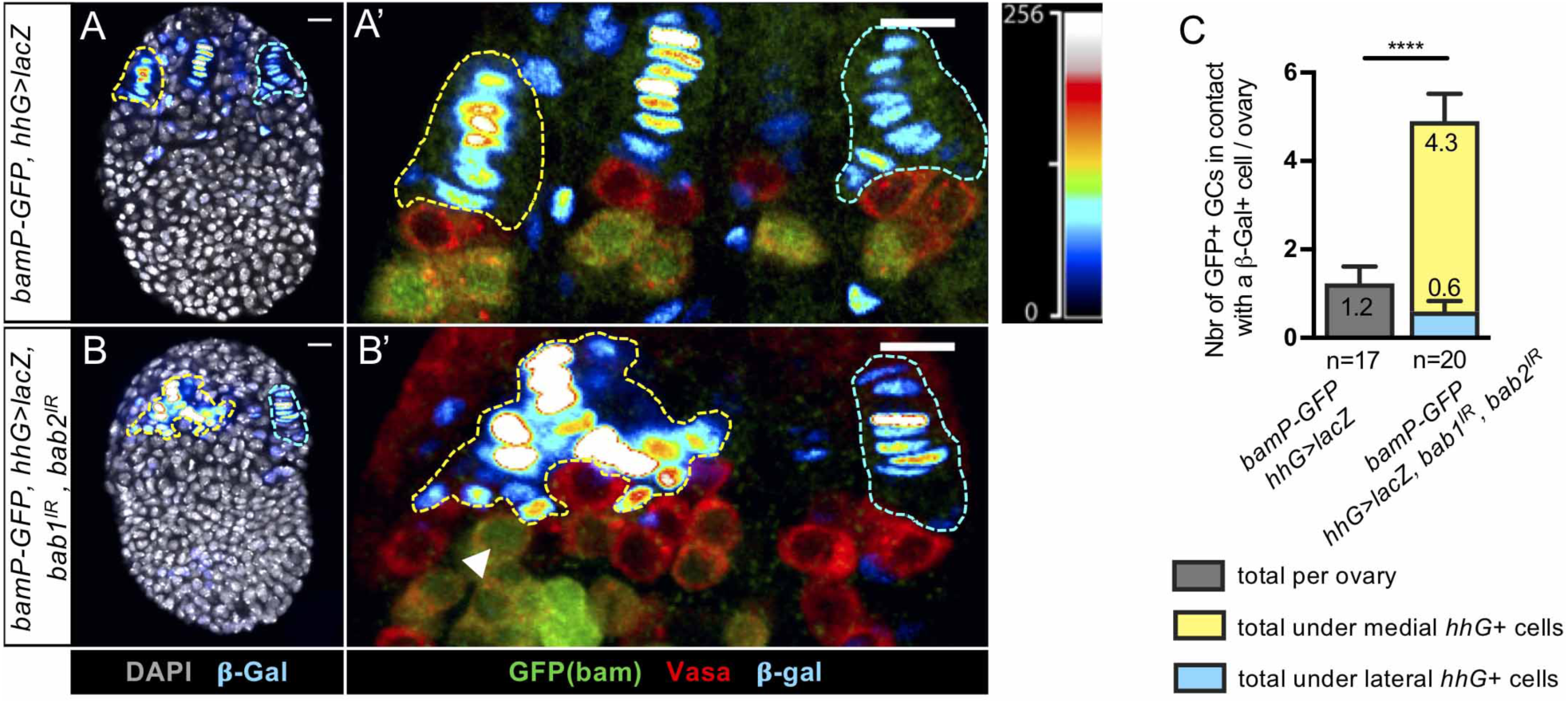
Some Germ Cells in contact with niche cells depleted of Bab proteins begin to differentiate into cystoblasts. (A-B’) Whole mount immunostaining of prepupal ovaries (projections of adjacent confocal sections) for detection of β-Galactosidase (β - Gal, Royal Lookup Table (LUT) Fiji, indicating signal intensity, inset to the right), GFP (green) and Vasa (red). Nuclei are labeled with DAPI (grey). Anterior is up, medial is left. Scale bars: 10 µm. *hhG*+ cells revealed by the detection of β-Gal (Royal LUT) are encircled in yellow and blue in the medial and lateral regions of the ovaries, respectively. (A’,B’) Higher magnification of the niche region of the corresponding ovaries in (A,B). Germ Cells (GCs) are marked with Vasa (red). (A’) In a control ovary (*bamP-GFP, hhG>lacZ),* the differentiating GCs expressing the GFP *bam* transcriptional reporter (green) are mainly found one-cell diameter away from the β-Gal+ (*hhG+*) cells (yellow and blue dotted lines). (B’) In ovaries depleted of Bab1 and Bab2, the differentiating GCs expressing the GFP *bam* transcriptional reporter (green) can also be observed in direct contact with β-Gal+ (*hhG+*) cells (white arrowhead). (C) Quantification of the number of GFP+ differentiating GCs in contact with β-Gal+ (*hhG+*) cells. In the ovaries depleted of Bab1 and Bab2, significantly more GFP+ differentiating GCs are found in contact with β-Gal+ (*hhG+*) cells and most of these cells are found in the medial part of the ovary where depletion of the Bab proteins is strong (yellow bar) rather than laterally where it is low (blue bar). Values are presented as means +s.e.m. p-values are calculated using a two-tailed t-test. n: sample size; ****: (p<0.0001).

### The functions of *bab1* and *bab2* are not required in niche cells for expression of Cap Cell specification markers

CCs have been shown to be essential for GSC establishment and maintenance (Panchal et al., 2017; Song et al., 2007; Ward et al., 2006; Xie and Spradling, 2000). Our results here show that Bab proteins are necessary in niche cells of the larval ovary for GSC establishment. One hypothesis is that *bab* function is necessary for specification of CCs which in turn would recruit the initial GSCs among PGCs. In order to test this possibility, we characterized the nature of the *hhG+* cells upon Bab1 and Bab2 depletion using a combination of markers that allow the distinction to be made between TF cells and CCs in prepupal niches: Traffic Jam (Tj) (Li et al., 2003), *P1444-LacZ* (Panchal et al., 2017), Delta (Panchal et al., 2017; Song et al., 2007) and *E(spl)mβ-CD2* (a transcriptional reporter of Notch activity) (de Celis et al., 1998).

The first two markers tested were Tj and *P1444-lacZ*. In prepupal control *hhG+* niches, *tj* and *P1444-LacZ* were expressed at high levels in CC nuclei (Fig 5A-5A’’, yellow bracket) and sometimes also in transition cells (the basal-most TF cell) (Fig 5A’-A’’’ and Fig 5C’, green arrowheads), at a low level for *tj*, consistent with what was previously reported (Panchal et al., 2017). *P1444-LacZ* was also expressed in some TF cells at a much lower level than in CCs and in transition cells (Fig 5A-5A’’, green bracket). Therefore, within niches, high levels of *tj* and *P1444-lacZ* expression designates CCs, while low levels or absence of *P1444-lacZ* expression without any Tj distinguishes TF cells. Upon depletion of Bab1 and Bab2 by RNAi specifically in niche cell precursors, the disorganized clustered medial *hhG+* cells were composed of two cell populations. One of the populations was posteriorly-positioned, adjacent to Vasa+ GCs, and presented Tj nuclear accumulation, comparable to that detected in control CCs (Fig 5B’ and 5B’’’ and Fig 5D’, yellow brackets), associated with the expression of *P1444-lacZ*, albeit at a lower level than in control CCs (Fig 5B’-5B’’, yellow bracket). Therefore, in posterior *hhG+* cells, these CC markers are present in the absence of Bab proteins. The second population of cells within the medial *hhG+* cell cluster positioned more anteriorly, like control TF cells, did not accumulate Tj (Fig 5B’ and 5D’, green brackets), but unlike control TF cells, did not express even low levels of *P1444-lacZ* (Fig 5B’-5B’’, green bracket). The identity of this second population may thus be mixed and this could be linked to the inability of these cells to form proper TFs.

**Fig 5.**
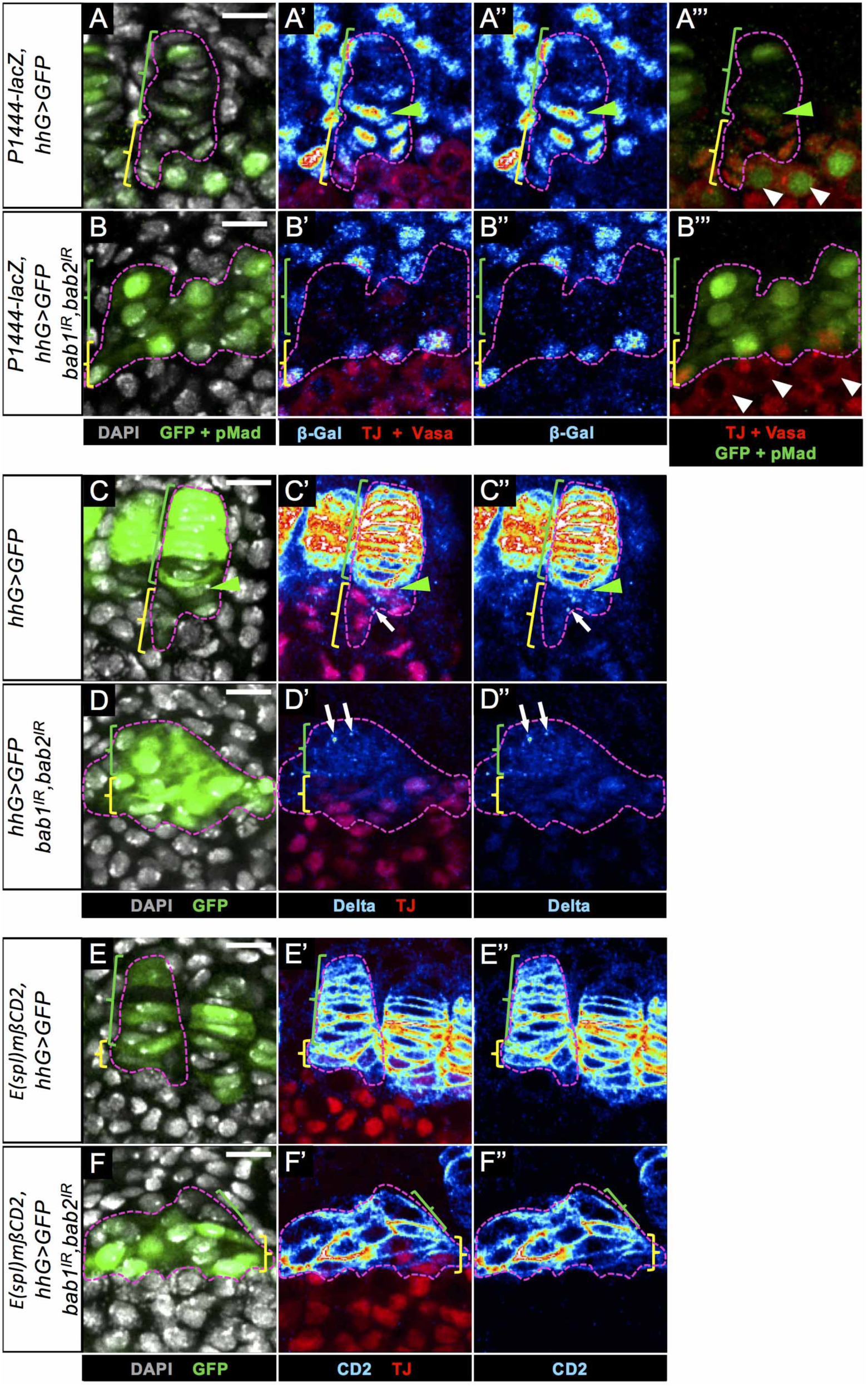
*bab1* and *bab2* functions in niche cells are not necessary for expression of several CC specification markers. (A-F) Whole mount immunostaining of the medial region of prepupal ovaries (projections of adjacent confocal sections). Nuclei are labelled with DAPI (grey). Anterior is up. Scale bars: 10 µm. (A-A’’’,C-C’’,E-E’’) correspond to the control ovaries. One *hhG>GFP* niche is encircled in each panel (pink dotted line). (B-B’’’,D-D’’,F-F’’) correspond to *bab1* and *bab2* RNAi-mediated knockdown ovaries. The entire cluster of *hhG+* cells depleted of Bab1 and Bab2 is encircled in each panel (pink dotted line). (A,C,E) The green and yellow brackets indicate Terminal Filament (TF) cells and Cap Cells (CCs), respectively, and the green arrowheads point to transition cells. (B,D,E) The green and yellow brackets indicate the anterior- and posterior-most *hhG+* cells, respectively. (A-A’’’) In the control ovaries, the two CCs markers, nuclear *P1444-lacZ* (Royal LUT) and nuclear Traffic Jam (red), are detected in CCs (yellow bracket) and in the transition cell (green arrowhead). Nuclear *P1444-lacZ* is also detected at a weaker level in some TF cells. The Germ Cells (GCs), marked with Vasa protein (red and cytoplasmic), in direct contact with niche cells (white arrowheads) show nuclear pMad (green) and are thus considered as Germinal Stem Cells (GSCs). (B-B’’’) Posterior-most medial *hhG+* cells depleted of Bab1 and Bab2, express both CC markers (Tj and *P1444-lacZ*) and are adjacent to GCs which do not present pMad+ (B’’’, arrowheads) and thus are not considered to be GSCs. (C-C”) In control ovaries, Delta (Royal LUT) accumulates at the plasma membrane and in cytoplasmic vesicles in TF cells, is detected in vesicle around the transition cell (green arrowhead) and sometimes in CCs (arrow). (D-D”) Upon depletion of Bab1 and Bab2 in *hhG+* cells, Delta is not present at the plasma membrane, but is found in some vesicles (arrows). (E-E”,F-F”) In both the control (E-E”) and upon depletion of Bab1 and Bab2 in *hhG+* cells (F-F”), the Notch pathway transcriptional reporter *E(spl)mβ-CD2* is expressed since CD2 (Royal LUT) accumulates at the plasma membrane of TF cells / anterior-most *hhG+* cells (green brackets) and of CCs / posterior-most *hhG+* cells (yellow brackets), also accumulating Tj.

We next tested for the presence of Notch pathway-associated niche markers, *i.e*. the Notch ligand Delta and a Notch transcriptional reporter, *E(spl)mβ-CD2*. It has been shown that during niche morphogenesis, CC specification depends on Notch signaling (Hsu and Drummond-Barbosa, 2011; Song et al., 2007). More recently, it was shown that the transition cell was associated primarily with vesicular Delta, designating an active Delta signal-sending state (Yatsenko and Shcherbata, 2018). Consequent activation of Notch signaling in adjacent ICs was proposed to allow their specification into CCs. In *hhG>GFP* control prepupal ovaries, TF cells presented high levels of Delta at the plasma membrane (Fig 5C-5C’’, green bracket), transition cells were associated with active Delta within cytoplasmic vesicles (Fig 5C-5C’’, green arrowhead), and CCs were largely devoid of Delta except in few vesicles (Fig 5C-5C’’, yellow bracket and white arrow), consistent with previous reports (Panchal et al., 2017; Yatsenko and Shcherbata, 2018). CD2 from the *E(spl)mβ-CD2* reporter was detected at the plasma membrane of both TF cells and CCs (Fig 5E-5E’’, green and yellow brackets, respectively), with highest levels in posterior-most TF cells. In anterior-most *hhG+* cells knocked down for *bab1* and *bab2*, Delta was detected only in its active cytoplasmic vesicular form (Fig 5D’ and 5D’’, green brackets and arrows), and not at the plasma membrane, unlike in control TF cells (compare to 5C’ and 5C”, green brackets), while posterior-most *hhG+*/Tj+ cells did not present any Delta at the plasma membrane nor in vesicles as for most of the control CC population (compare yellow brackets in Fig 5D’ and 5D” to Fig 5C’ and 5C”). Despite this, all *hhG+* cells depleted of Bab1 and Bab2 expressed *E(spl)mβ-CD2* reporter indicative of active Notch signaling (Fig 5F’ and 5F’’, green and yellow brackets), as observed in control niche cells. Therefore, depletion of Bab proteins in niche cells did not prevent Notch signaling activation in these cells. This is likely due to vesicular Delta present in the more anterior Tj-/*hhG+* cells, since at the end of the third instar larval stage, the most posterior TF cells become Delta-sending cells presenting predominantly vesicular Delta and activating Notch signaling in adjacent CC precursors (Yatsenko and Shcherbata, 2018). Altogether, these results show that *bab1* and *bab2* are not necessary for several aspects of CC specification during niche formation since posterior-most *hhG+* cells depleted of Bab proteins present the appropriate expression pattern for all CC markers tested. Despite the presence of CC-marker expressing cells upon Bab depletion in niches, Vasa+ GCs in contact with these CCs are not GSCs since they are not activated for BMP signaling as evidenced by the absence of nuclear pMad (Fig 5B’’’ compare to Fig 5A”’, arrowheads). Therefore, the absence of GSC establishment with Bab-depleted *hhG+* cells cannot be attributed to failure in CC specification as far as the markers tested here.

### *bab1* and *bab2* are necessary for *dpp* transcription in niche cells at the time of initial GSC establishment

Since Bab proteins do not seem to be necessary for CC specification in the larval ovary, the absence of GSC establishment at this stage upon depletion of these transcription factors in niche cells could be due to a specific defect in *dpp* expression in CCs. Therefore, we tested whether the *bab* locus function controls *dpp* expression in CCs. For this we used two lines of the GFP-tagged FlyFos TransgeneOme library (Sarov et al., 2016), hereafter called *dpp-GFP* and *dpp-nlsGFP*. Each line contains a construction with a large genomic region (about 44 kb) covering both the *dpp* regulatory and coding sequences into which a C-terminal GFP tag was introduced. We first characterized the adult ovarian expression profile of these two lines. In adult germaria (Fig S1), we found that the two fusion proteins were detected at high levels in CCs (S1A’’’,B’’’ Fig, yellow brackets), sometimes in ECs (S1B Fig, blue brackets) and absent from TF cells (S1A’’’,B’’’ Fig, green brackets), and were also detected in prefollicular cells (S1B Fig, orange brackets), reflecting the endogenous *dpp* expression pattern (Liu et al., 2010, 2015; Luo et al., 2017; Wang et al., 2008; Xie and Spradling, 2000). For analysis in prepupal ovaries, we used the *dpp-nlsGFP* transgene since the nuclear Dpp:GFP fusion protein produced was less likely to perturb endogenous Dpp function. Indeed, no obvious alteration of wild type phenotype was observed in ovaries of females carrying this transgene (Fig 6A). At the prepupal stage, in control medial niches, this fusion protein was detected in CCs, and sometimes in transition cells, but not in other TF cells (Fig 6A-6A’’, yellow bracket, green arrowhead, and green bracket, respectively). In striking contrast, upon depletion of Bab1 and Bab2 in *hhG+* cells, *dpp-nlsGFP* was not expressed in the posterior-most *hhG+* cells positive for Tj and located next to GCs in the medial region of the ovary, that we have characterized as CCs (Fig 6B-6B’’, yellow bracket).

**Fig 6.**
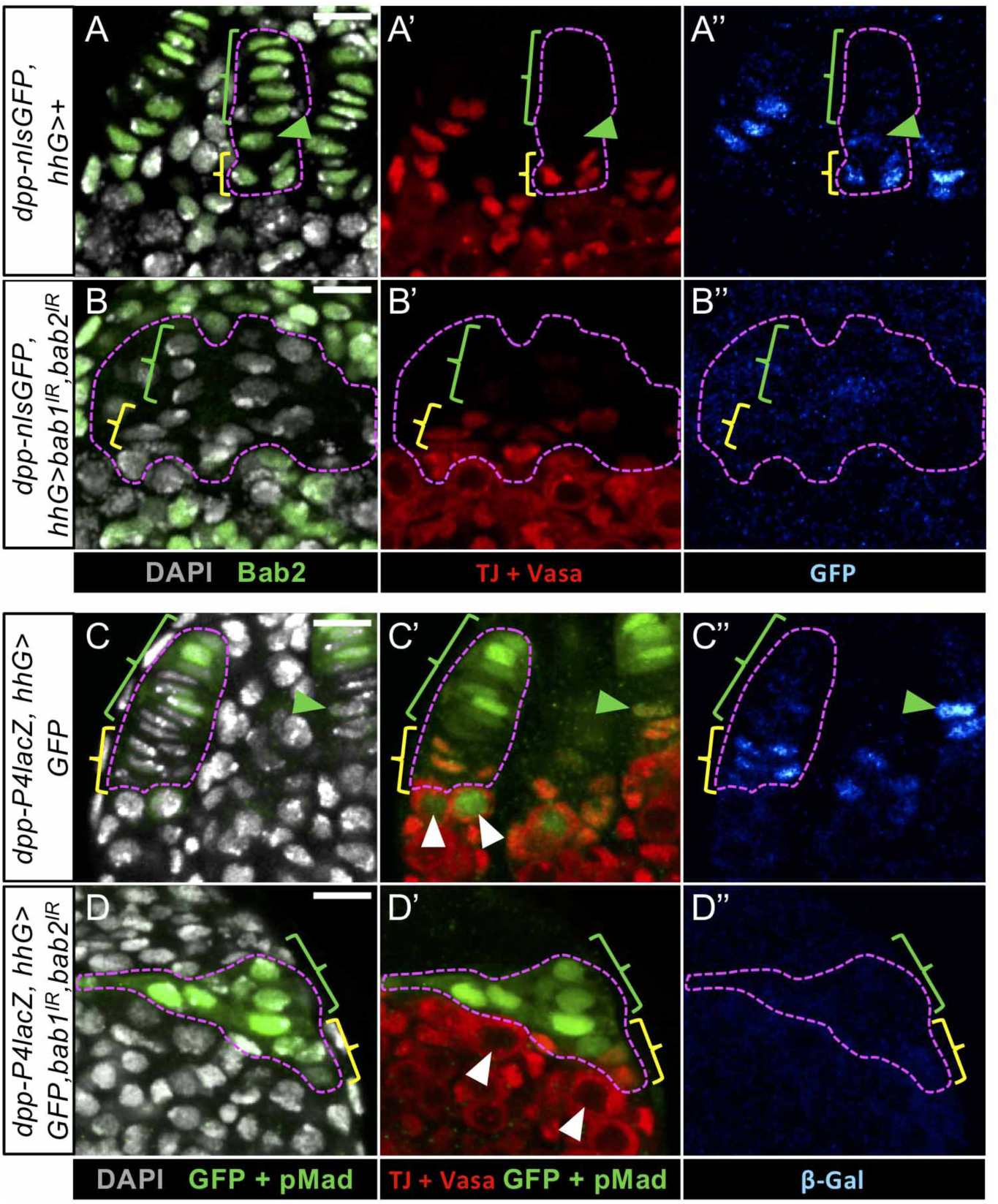
*bab1* and *bab2* functions in niche cells are required for *dpp* expression. (A-D) Whole mount immunostaining of the medial region of prepupal ovaries (projections of adjacent confocal sections). Nuclei are labeled with DAPI (grey). Anterior is up. Scale bars: 10 µm. (A’-D’) GCs are visualized with cytoplasmic Vasa (red). One *hhG>GFP* niche is encircled in images of control ovaries (A-A”,C-C”) and green and yellow brackets indicate Terminal Filament (TF) cells and Cap Cells (CCs), respectively. The green arrowheads point to transition cells. (B-B”, D-D”) The entire cluster of medial *hhG+* cells is encircled in images of *bab1* and *bab2* knockdown ovaries, and green and yellow brackets indicate the anterior- and posterior-most *hhG+* cells, respectively. (A,B) Somatic cells are marked with Bab2 (green). (A-A”) In a control ovary (*dpp-nlsGFP*, *hhG>+), dpp-nlsGFP* (Royal LUT) is specifically expressed in the Cap Cells (CCs) (yellow bracket) along with nuclear Traffic Jam (Tj, red) and sometimes in the transition cell (green arrowhead), but not in Terminal Filaments (green bracket). (B-B”) Upon *bab1* and *bab2* RNAi-mediated knockdown (*dpp-nlsGFP*, *hhG>bab1^IR^,bab2^IR^), dpp-nlsGFP* is not expressed in posterior-most *hhG+* cells (yellow brackets) that are positive for the nuclear CC marker Tj (red), and in contact with the Germ Cells (GCs) marked with cytoplasmic Vasa (red). (C-C”) In a control ovary (*dpp-P4lacZ, hhG>GFP*), *dpp-P4lacZ* expression is found in CCs (yellow brackets) in contact with GSCs marked with cytoplasmic Vasa (red) and nuclear pMad+ (green) (white arrowheads) and often in the transition cell (green arrowhead). (D-D’’) Upon *bab1* and *bab2* RNAi-mediated knockdown (*dpp-P4lacZ*, *hhG>bab1^IR^,bab2^IR^)*, *dpp-P4lacZ* expression is not detected in the posterior-most *hhG+* cells, positive for the CC marker Tj (yellow brackets), which correlates with an absence of pMad in the Vasa+ underlying GCs (white arrowheads).

Another *dpp* transcriptional reporter, *dppP4-lacZ*, contains a much smaller region of the cis-regulatory sequences present in *dpp-nlsGFP*. *dppP4-lacZ* was shown to be expressed in CCs as endogenous *dpp*, but also in TF cells in adult wild-type niches (Li et al., 2016), thereby not fully recapitulating the endogenous *dpp* expression pattern. We found here that in the medial region of prepupal control ovaries, this reporter, like *dpp-nlsGFP* and endogenous *dpp* (Matsuoka et al., 2013), was expressed in CCs (Fig 6C-6C’’, yellow bracket) and often in transition cells (Fig 6C-6C’’, green arrowhead), but not in other TF cells (Fig 6C-6C’’, green bracket). These *dppP4-lacZ+* CCs were in contact with pMad+/Vasa+ GSCs (Fig 6C’, white arrowheads). In medial *hhG+* cells depleted of Bab1 and Bab2, *dppP4-lacZ* was not or very faintly expressed (Fig 6D-6D’’) and this correlated with the absence of pMad in adjacent GCs (Fig 6D’, white arrowheads). Altogether, these results indicate that Bab1 and Bab2 are necessary for *dpp* transcription in CCs at the prepupal stage. In addition, since *hhG+* cells depleted of Bab proteins express some CC markers but do not express *dpp* transcriptional reporters, they must thus be considered only as “CC-like”.

### *bab* regulates Engrailed/Invected accumulation in TF cells, but not in CCs, during larval stages

Like the *bab* genes, *engrailed* and its paralog *invected (en/inv)* have also been shown to be involved in proper TF formation (Bolívar et al., 2006). In addition, evidence strongly suggests that En directly controls *dpp* expression in adult CCs (Luo et al., 2017). Therefore, we addressed whether *bab* function may act upstream of *en/inv* in larval stages, and thereby would impact *dpp* transcriptional regulation indirectly through *en/inv*. In order to explore possible functional interactions between Bab and En/Inv proteins, we tested whether *en/inv* expression in niche cells of prepupal ovaries depends on Bab proteins. It has been reported that *en*/*inv* are specifically expressed in GSC niche cells as of the beginning of TF formation in the larval ovary (Allbee et al., 2018; Forbes et al., 1996; Green and Extavour, 2012; Mendes and Mirth, 2016). In addition, we found here a significant difference in En/Inv levels between TF cells and CCs in control prepupal niches (*hs-FLP; FRT-bab^AR07^*/*FRT-GFP* and *hhG>GFP,* Fig 7A and 7A’ and Fig 7B and 7B’, green and yellow brackets, respectively), with a three-fold higher level in control TF cells than CCs (*hs-FLP; GFP*, Fig 7D). Induction of mitotic TF cell clones homozygous for *bab^AR07^*, a null allele for both *bab1* and *bab2* (Couderc et al., 2002), and marked by absence of GFP expression (Fig 7A, green arrowhead) showed that En/Inv levels were reduced by two-fold, cell autonomously in *bab^AR07^* TF cell clones (*hs-FLP; bab^AR07^*) when compared to control TF cells (*hs-FLP; GFP*) in the same ovaries (Fig 7A and 7A’, green arrowhead compared to green bracket, and Fig 7D for quantification). In contrast, En/Inv levels in *bab^AR07^* CC clones (Fig 7A and 7A’, yellow arrowheads) were not significantly different from wild-type CCs in the same ovaries (Fig 7A and 7A’, yellow arrowheads compared to yellow bracket, and Fig 7D for quantification).

**Fig 7.**
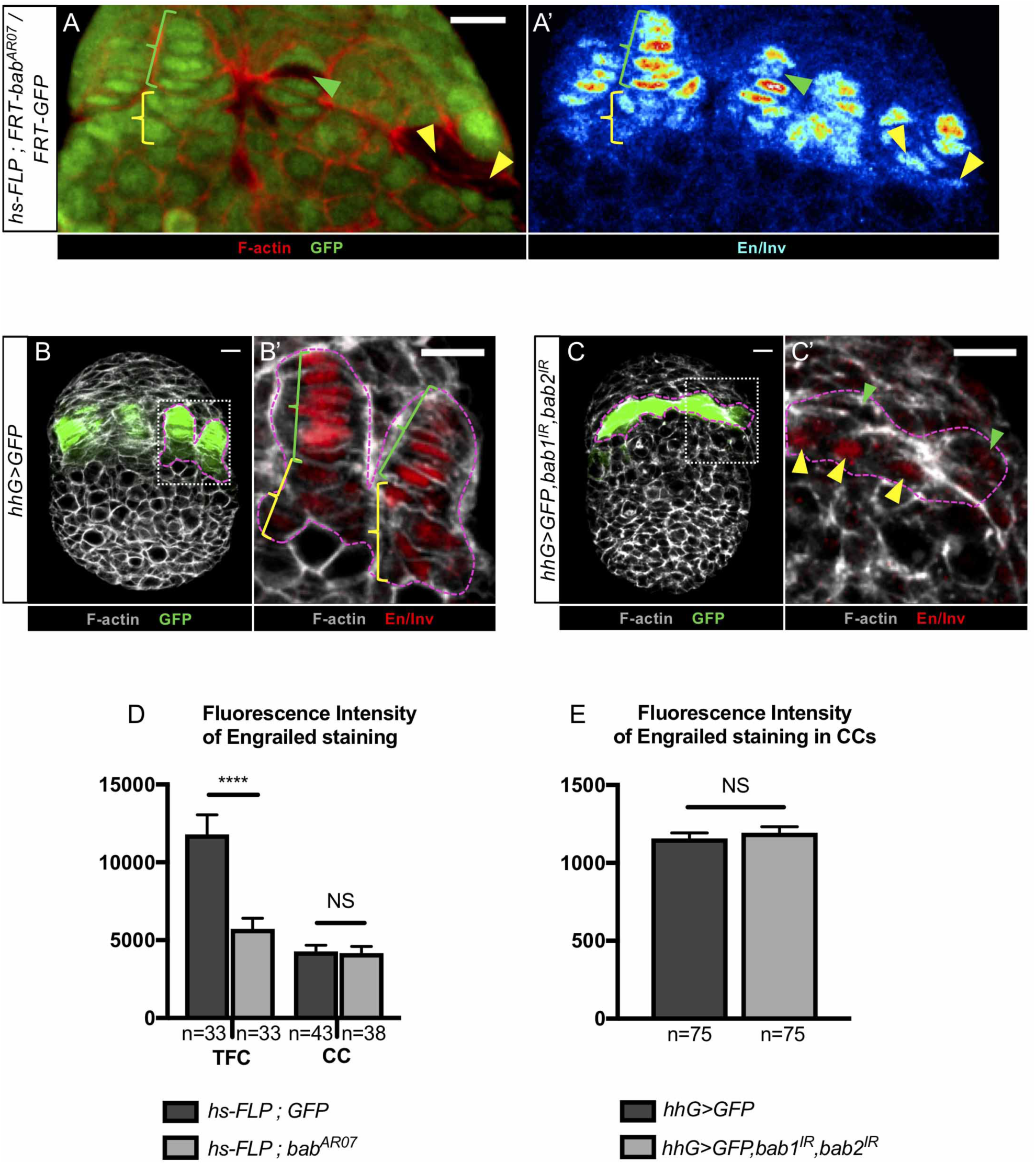
*bab* locus regulates Engrailed/Invected accumulation in Terminal Filament cells but not in Cap Cells. (A-C’) Whole mount immunostaining of prepupal ovaries (projections of adjacent confocal sections) for detection of GFP (green) and En/Inv shown in Royal LUT (A’) and red (B’ and C’). F-actin labeling is shown in red (A) and grey (B-C’). Anterior is up, medial is left. Scale bars: 10 µm. (A, A’, B’) Green and yellow brackets indicate Terminal Filament (TF) cells and Cap Cells (CCs), respectively. (C,C’) The entire cluster of medial *hhG+* cells depleted for Bab1 and Bab2 is encircled, and green and yellow arrowheads indicate the anterior- and posterior-most *hhG+* cells, respectively. (A,A’) Niche region of a mosaic ovary (*hs-FLP; FRT-bab^AR07^/FRT-GFP),* containing mitotic cell clones homozygous for the *bab^AR07^* mutation and marked by absence of GFP (green and yellow arrowheads). In *bab^AR07^* mutant Terminal Filament (TF) cells (green arrowheads), the signal for Engrailed/Invected (En/Inv) is lower than that in wild type TF cells (green brackets). However, in *bab^AR07^* mutant Cap Cells (CCs) (yellow arrowheads), the level of En/Inv is similar to its endogenous level in wild type CCs (yellow bracket). (B,C) Prepupal ovaries expressing GFP in the niche cells (*hhG>GFP)*, as well as RNAi against *bab1* and *bab2 (hhG>GFP, bab1^IR^, bab2^IR^*^)^ in C). (B’,C’) Higher magnification of the regions framed with dotted lines in the corresponding ovaries. The knockdown of *bab1* and *bab2* in TF cells and CCs leads to the presence of anterior-most cells *hhG+* cells (C’, green arrowheads) with lower levels of En/Inv compared to TF cells in the control (B’, green brackets). In contrast, the posterior-most *hhG+* cells depleted of Bab1 and Bab2 (C’, yellow arrowheads), show a similar level of En/Inv compared to that in control CCs (B’, yellow brackets). (D) Graph comparing the fluorescence intensity of En/Inv in control TF cells and CCs compared to the corresponding *bab^AR07^* mitotic clonal cells (A). In *bab^AR07^* mutant TF cells (*hs-FLP*, *bab^AR07^*), the En/Inv fluorescence intensity is 2-fold lower than in adjacent control TF cells (*hs-FLP*, *GFP*). However, in *bab^AR07^* mutant CCs, the En/Inv fluorescence intensity is similar to that in the adjacent control CCs. (E) Graph comparing the fluorescence intensity of En/Inv in CCs in the control (B’, yellow brackets) and in posterior-most *hhG+* cells (C’, yellow arrowheads) upon *bab1* and *bab2* RNAi-mediated knockdown. There is no significant difference between En/Inv levels in the two genetic contexts. Values are presented as means +s.e.m. p-values are calculated using a two-tailed t-test. n: sample size; NS: Not Significant (p>0.05); ****: p<0.0001).

Using RNAi to knockdown *bab* locus function in niche cells (*hhG>GFP, bab1^IR^, bab2^IR^* ovaries), similar results were obtained, with En/Inv levels being significantly lower in anterior-most *hhG+* cells depleted for Bab1 and Bab2 (Fig 7C’, green arrowheads) than in control TFs (Fig 7B’, green brackets), while present at the same levels in posterior-most *hhG+* cells depleted for Bab1 and Bab2 in contact with GCs (Fig 7C’, yellow arrowheads) than in control CCs (Fig 7B’, yellow brackets, and Fig 7E for quantification). Taken together these results show that *bab* function is required cell-autonomously in TF cells of larval ovaries to ensure high En/Inv accumulation in these cells. On the contrary, *bab* function is not required in CCs to ensure normal level of En/Inv. Therefore, the function of Bab proteins for normal *dpp* expression in CCs cannot be linked to their control of En/Inv accumulation in these cells.

### Depletion of *Engrailed/Invected* in niches does not prevent either TF formation or initial GSC establishment

As shown above, strong depletion of Bab proteins in TF cells led to both lower accumulation of En/Inv in these cells and highly defective TF morphogenesis. Moreover, elimination of En/Inv by induction of mitotic TF cell clones homozygous for the *en^E^/inv^E^* null mutant allele, affecting both paralogs, was previously shown to lead to a mild defect in alignment of TF cells in larval ovaries, which was aggravated by pupal stages (Bolívar et al., 2006). We thus tested whether depletion of En/Inv in all niche cells would phenocopy the *bab* loss of function strong TF morphogenesis defect. To achieve strong depletion of *en/inv* in niches cells, we used a *bab-Gal4* driver (hereafter named *babG*), expressed in all ovarian somatic cells including niche cells (Cabrera et al., 2002) and Gal80^TS^ which allows Gal4 activity at the Gal80^TS^ restrictive temperature (29°C) and impedes Gal4 activity at the Gal80^TS^ permissive temperature (18°C). When the *babG* driver was expressed during all larval development, very strong En/Inv depletion was obtained in prepupal ovary niches (Fig 8B and Fig 9B,B’) compared to controls (Fig 8A and Fig 9A,A’). Under these conditions, normally formed TFs were present (Fig 8B-8B” compared to control in 8A-8A’’), and the number of TFs per ovary (Fig 8C), as well as the number of TF cells per TF (Fig 8D) and of CCs per TF (Fig 8E) were not significantly different from controls. These results indicate that *en/inv* function is not required for individual TF formation. Thus, even though *bab* function is required for accumulation of high levels of En/Inv in TF cells, the function of *bab* in TF formation most likely does not depend on *en/inv* function since these two genes are not required in this process.

**Fig 8.**
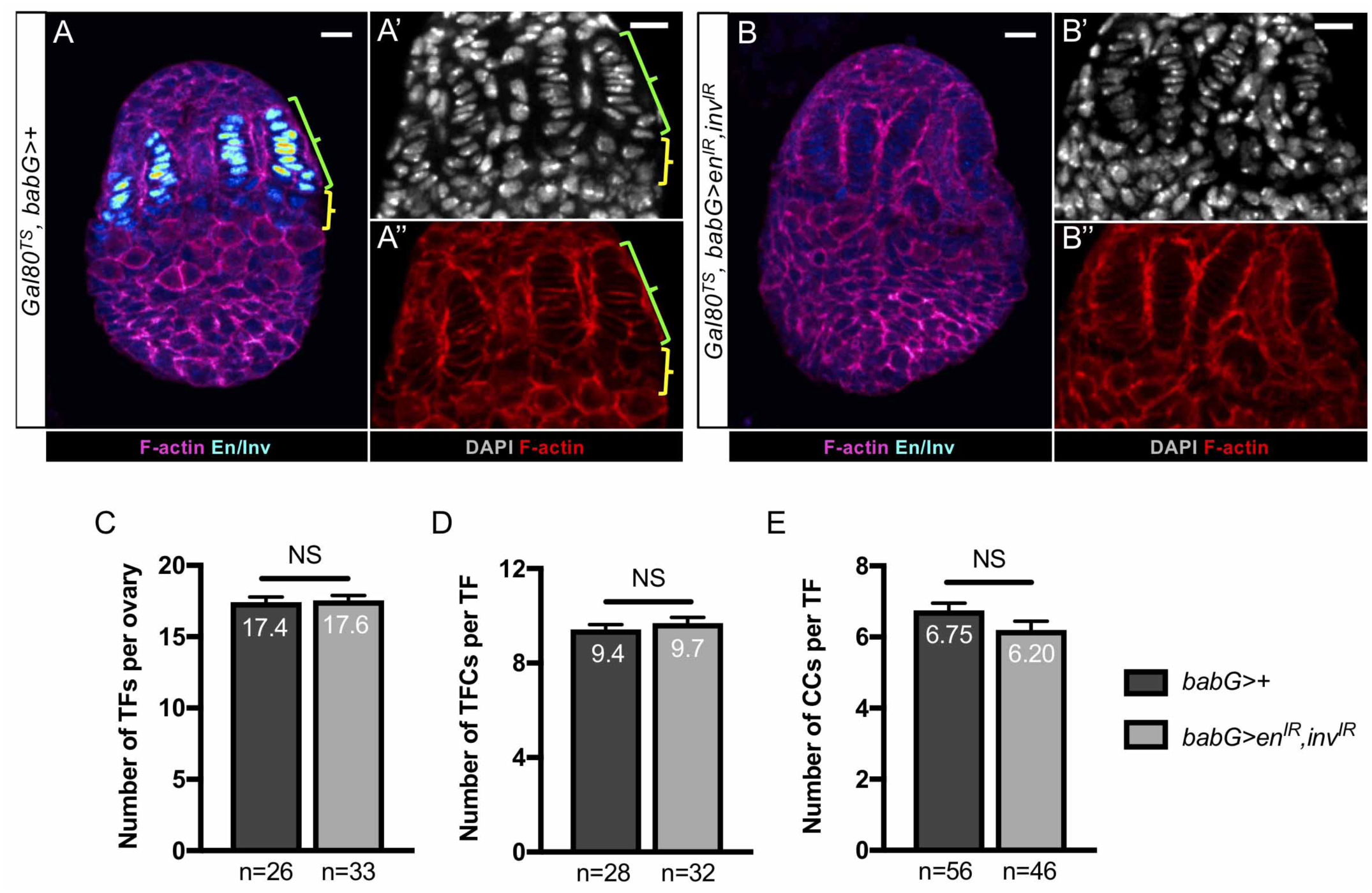
TF formation does not rely on *en/inv* function. (A-B) Whole mount immunostaining of prepupal ovaries (projections of adjacent confocal sections) for detection of Engrailed/Invected (En/Inv) (Royal LUT). Nuclei are labeled with DAPI (grey). F-actin labeling is shown in magenta and red (A,B and A”,B”, respectively). Anterior is up, medial is left. Scale bars: 10 µm. (A’,A’’,B’,B’’) Higher magnifications of the niche regions of the corresponding ovaries in (A,B). (A-A’’) In a control ovary (*Gal80^TS^*, *babG>*+) En/Inv accumulate in the niche cells: at high levels in Terminal Filament cells (TF cells; green bracket) and at lower levels in Cap Cells (CCs; yellow bracket). (B-B’’) In an ovary carrying transgenes targeting *en/inv* for RNAi (*babG/UAS-en^IR^, UAS-inv^IR^*) these two proteins are efficiently depleted (B), while TFs are well formed (B”) and TF cells properly aligned within TFs (B’). (C-E) Graphs comparing different parameters related to TF formation, between control ovaries and ovaries depleted of En/Inv. (C) number of TFs per ovary, (D) number of TF cells per TF and (E) number of CCs per TF. The TF cells (green brackets in A-A”) were distinguished from CCs (yellow brackets in A-A”) by their flattened nuclei and their flat shape, as determined by DAPI (grey, A’ and B’) and F-actin labeling (red, A” and B”), respectively. CCs were distinguished from Intermingled Cells (ICs) using Bab1 immunostaining, which is high in CCs and low in ICs (data not shown) and from TF cells due to the round shape of their nuclei. None of these parameters were significantly different between control and En/Inv depleted ovaries. Values are presented as means +s.e.m. p-values are calculated using a two-tailed t-test. n: sample size; NS: Not Significant (p>0.05).

**Fig 9.**
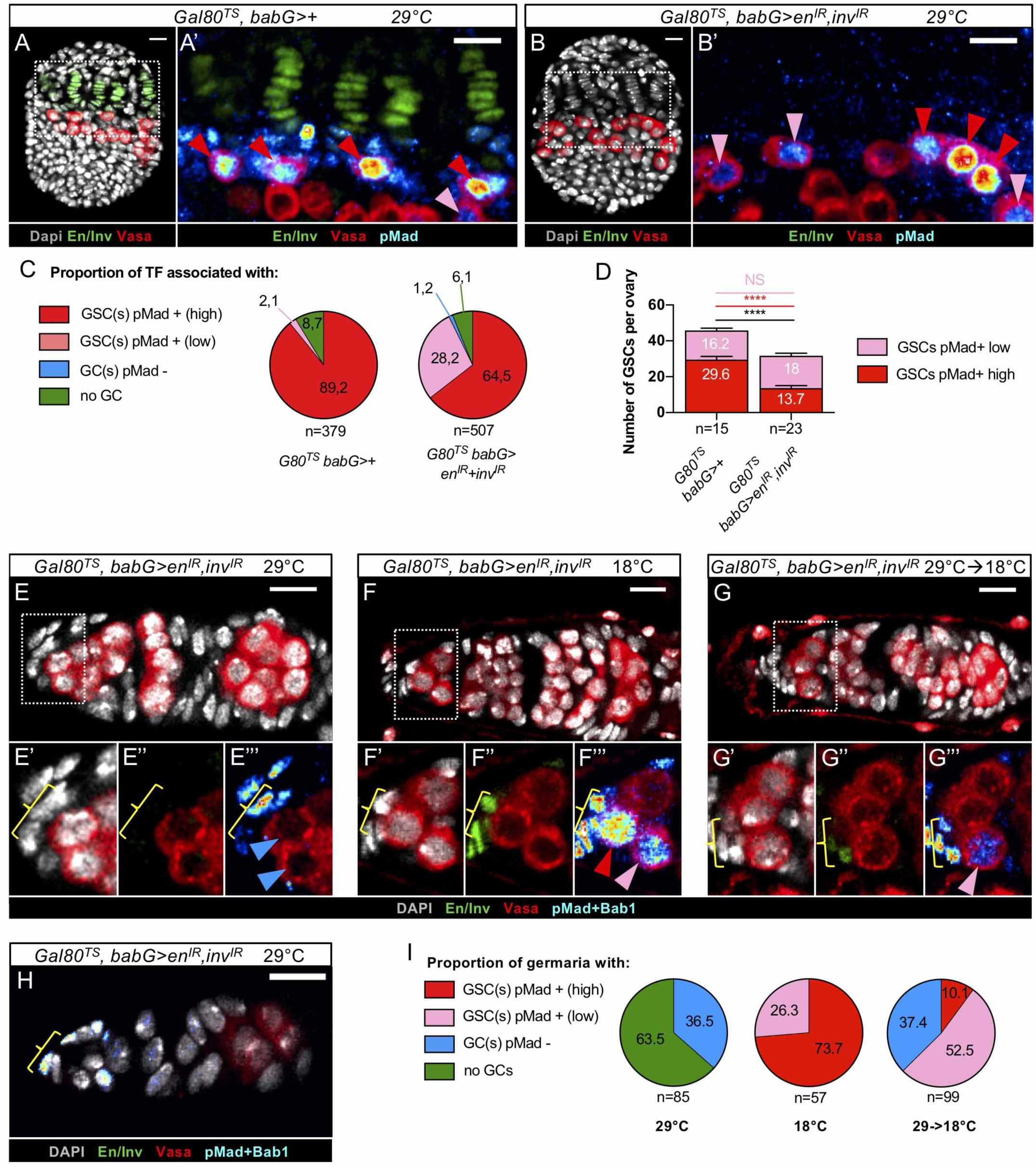
En/Inv are not necessary for GSC establishment but are involved in their maintenance. (A-B) Whole mount immunostaining of prepupal ovaries (projections of adjacent confocal sections) for detection of Engrailed/Invected (En/Inv) (green) and Vasa (Royal LUT). Nuclei are labeled with DAPI (grey). Anterior is up, medial is left. Scale bars: 10 µm. (A’,B’) Higher magnifications of the niche regions of the corresponding ovaries in (A,B). In control ovaries (*Gal80^TS^; babG>+*) and ovaries depleted of En/Inv *(Gal80^TS^; babG/UAS-en^IR^, UAS-inv^IR^),* Germinal Stem Cells (GSCs) with both high and low pMad levels are present (red and pink arrowheads, respectively). (C) Pie chart comparing the proportion of prepupal niches associated with: at least one GSC with a high level of pMad (red), at least one GSC with a low level of pMad (pink), GCs without nuclear pMad (blue) and no Germinal Cells (GC) (green). The depletion of En/Inv does not change the proportion of Terminal Filaments (TFs) associated with at least one GSC (sum of pink and red portions). (D) Graph comparing the number of GSCs per prepupal ovary in control and En/Inv depleted ovaries. The number of GSCs per ovary is significantly lower in En/Inv depleted ovaries (black bar), due to the significantly lower number of high-level pMad+ GSCs (red bar) and not to the number of low-level pMad+ GSCs (pink bar) which are not significantly different. Values are presented as means +s.e.m. p-values are calculated using a two-tailed t-test. n: sample size; NS: Not Significant (p>0.05); ****: (p<0.0001). (E-H) Germaria from 1 day-old adult females carrying transgenes for temperature-controlled RNAi of *en/inv* (*Gal80^TS^; babG/UAS-en^IR^, UAS-inv^IR^),* immunostained for En/Inv (green), pMad and Bab1 (Royal LUT) and Vasa (red). Nuclei are labelled with DAPI. Images are projection of confocal sections. Anterior is up and the scale bars represent 10µm. The transgenes present and temperatures used for raising the flies are indicated on top of the images. (E’-E’’’,F’-F’’’, G’-G’’’) Higher magnifications of the corresponding niche regions in (E,F,G). (E-E”’) Germarium of a female raised at 29°C from the L1 stage to dissection shows efficient depletion of En/Inv in Cap Cells (CCs) (9E’’, yellow bracket). This germaria does not contain any pMad+ GSCs (E’’’, blue arrowheads). (F-F”’) Germarium of a female raised at 18°C from the L1 stage to dissection is not depleted of En/Inv in CCs (F’’, yellow bracket) and exhibits GSCs with high and low levels of pMad (F’’’, red and pink arrowheads, respectively). (G-G”’) Germarium of a female raised at 29°C from L1 stage and transferred to 18°C at the prepupal stage, showing the presence of En/Inv in CCs (G’’, yellow bracket) and a pMad+ GSCs (G’’’, pink arrowhead). (H) Germarium of a female raised at 29°C from the L1 stage to dissection showing efficient depletion of En/Inv in CCs (yellow bracket). This germaria does not contain any GCs (Vasa, red) close to the niche. (I) Pie chart comparing the proportion of germaria with: at least one GSC with a high level of pMad (red), at least one GSC with a low level of pMad (pink), GCs without nuclear pMad (blue), or no GCs (green). The re-expression of en*/inv* in ovaries from the prepupal stage onwards (29->18°C) leads to partial rescue (from 0% to 62.6%) of the proportion of germaria containing GSCs.

In adult ovary GSC niches, *en/inv* is known to be necessary for GSC maintenance through its regulation of *dpp* expression in CCs likely by direct binding of En to *dpp* regulatory sequences (Luo et al., 2017; Rojas-Ríos et al., 2012). We tested whether *en/inv* gene functions are also necessary for GSC establishment in ovaries by the prepupal stage. Both control and En/Inv-depleted ovaries displayed niches with high and low pMad+ GSCs (Fig 9A and 9A’ and Fig 9B and 9B’, red and pink arrowheads, respectively) and the proportion of TFs associated with GSCs was similarly high in both contexts (91.3% and 92.7%, respectively, Fig 9C). The average number of GSCs per ovary, however, was about 30% lower in ovaries depleted of En/Inv when compared to control ovaries (Fig 9D). This difference corresponds to a lower proportion of high pMad+ GSCs per ovary in the *en/inv* RNAi context (about half as many, 29.6 vs. 13.7 on average, Fig 9D). Therefore, En/Inv-depleted niches are almost all associated with GSCs but they contain fewer GSCs per niche than control niches and in particular fewer high-pMad+ GSCs per niche (Fig 9C). It is possible that En/Inv-depleted niches initially recruit GSCs with high pMad levels normally during niche formation, but that these niches do not have the capacity to maintain GSCs efficiently to the prepupal stage that was analyzed. In order to address this possibility, we tested the fate of GSCs, recruited initially within larval En/Inv-depleted niches, in young adult female siblings of the same genotype (*Gal80^TS^, babG> en^IR^, inv*^IR^). GSC niches of these 1 day-old female siblings maintained at 29°C as of the prepupal stage (Fig 9E-9E’’’) were efficiently depleted of En/Inv (Fig 9E-9E’’’), compared to niches from control females (Fig 9F-9F”’) always maintained at 18°C (Fig 9F-9F”’, yellow brackets indicate CCs). En/Inv depletion was associated with 100% of germaria lacking pMad+ GSCs (Fig 9E’’’, blue arrowheads compared to control in 9F’’’, red and pink arrowheads, and Fig 9I), with a majority of germaria even devoid of any Vasa+ GCs (Fig 9H and 9I). This result indicates that all newly established GSCs in each niche of prepupal ovaries depleted of En/Inv (Fig 9B’, red and pink arrowheads) were no longer present at adulthood (Fig 9E’’’, blue arrowheads). This result supports the hypothesis that in the absence of En/Inv, GSCs are established at larval stages but that they are not maintained during pupal stages. This is supported by the fact that partial re-expression of *en/inv* in niche cells from the onset of pupariation (29°C to 18°C switch, Fig 9G-9G’’’, yellow bracket) substantially rescued the adult GSC loss phenotype (Fig 9G-9G’’’, pink arrowhead) from 0% to more than 60% (Fig 9I). Altogether, these results strongly suggest that *en/inv* functions are not essential for initial GSC establishment within newly formed niches during larval stages. However, they are essential for GSC maintenance in niches starting from at least the beginning of the pupal stage, but possibly even earlier, as soon as GSCs are established.

### Overexpression of *bab2* in germaria leads to GSC-like tumors

Since Bab proteins are necessary for TF morphogenesis and initial GSC establishment, we then asked whether these proteins could be sufficient to induce formation of ectopic TFs and/or GSCs. With this aim, we expressed *UAS-bab1* or *UAS-bab2* constructs with the *C587-Gal4* driver (called *C587G* hereafter) at either 25°C or 29°C. *C587G* is expressed in most, but not all, somatic cells of developing female gonads (Zhu and Xie, 2003). With these tools, we generated prepupal ovaries with higher levels of Bab1 in ICs than in control ovaries (S2B Fig, arrowheads point to several ICs, compare to S2A Fig, bracket) and ectopic Bab1 in more basal somatic cells. For Bab2, higher levels were particularly obtained in ICs compared to the control (S2C’ Fig, arrowheads point to ICs, compare to S2A’, bracket). We found that ICs with higher Bab1 or Bab2 levels than normal did not exhibit En/Inv accumulation just as control ICs (S2B”and S2C” Fig, arrowheads compare to S2A”, bracket). In addition, we did not detect any obvious change in Bab1 levels in cells overexpressing *bab*2, nor any change in Bab2 levels in cells ectopically expressing Bab1 (S2B’ Fig compared to S2A’ Fig and S2C Fig compared to S2A Fig). Finally, none of these conditions appeared to affect the organization of pupal ovaries (S2 Fig).

In adult germaria, *C587G* has been shown to be expressed only in ECs and early prefollicular cells (Song et al., 2004). In order to test the effect of expressing *bab1* ectopically or over-expressing *bab2* in adult ovaries, we analyzed ovaries from 10-day old females raise at 29°C as of eclosion. Under these conditions, we detected ectopic Bab1 in ECs and prefollicular cells near the middle of germaria (Fig 10B,B’ compared to 10A,A’, blue and yellow arrowheads, respectively). As for Bab2, significantly higher levels were induced in both ECs and prefollicular cells than in the control (Fig 10C-10C” compared to 10A-10A”, blue and yellow arrowheads, respectively). The levels of Bab1 and Bab2 accumulation in ECs and prefollicular cells using UAS transgenes did not appear different from endogenous levels of these two proteins in niche cells of the same germaria (Fig 10B’ and 10C’’, blue and yellow arrowheads point to ECs and prefollicular cells, respectively), compared to yellow brackets (CCs)). In ECs and prefollicular cells ectopically expressing *bab1* or overexpressing *bab2*, no cross-regulation between the two genes was apparent (Fig 10B’,B’’ and Fig 10C’,C”, yellow arrowheads). Moreover, abnormal accumulation of Bab1 and Bab2 in ECs and prefollicular cells did not lead to any obvious morphological changes of these cells.

**Fig 10.**
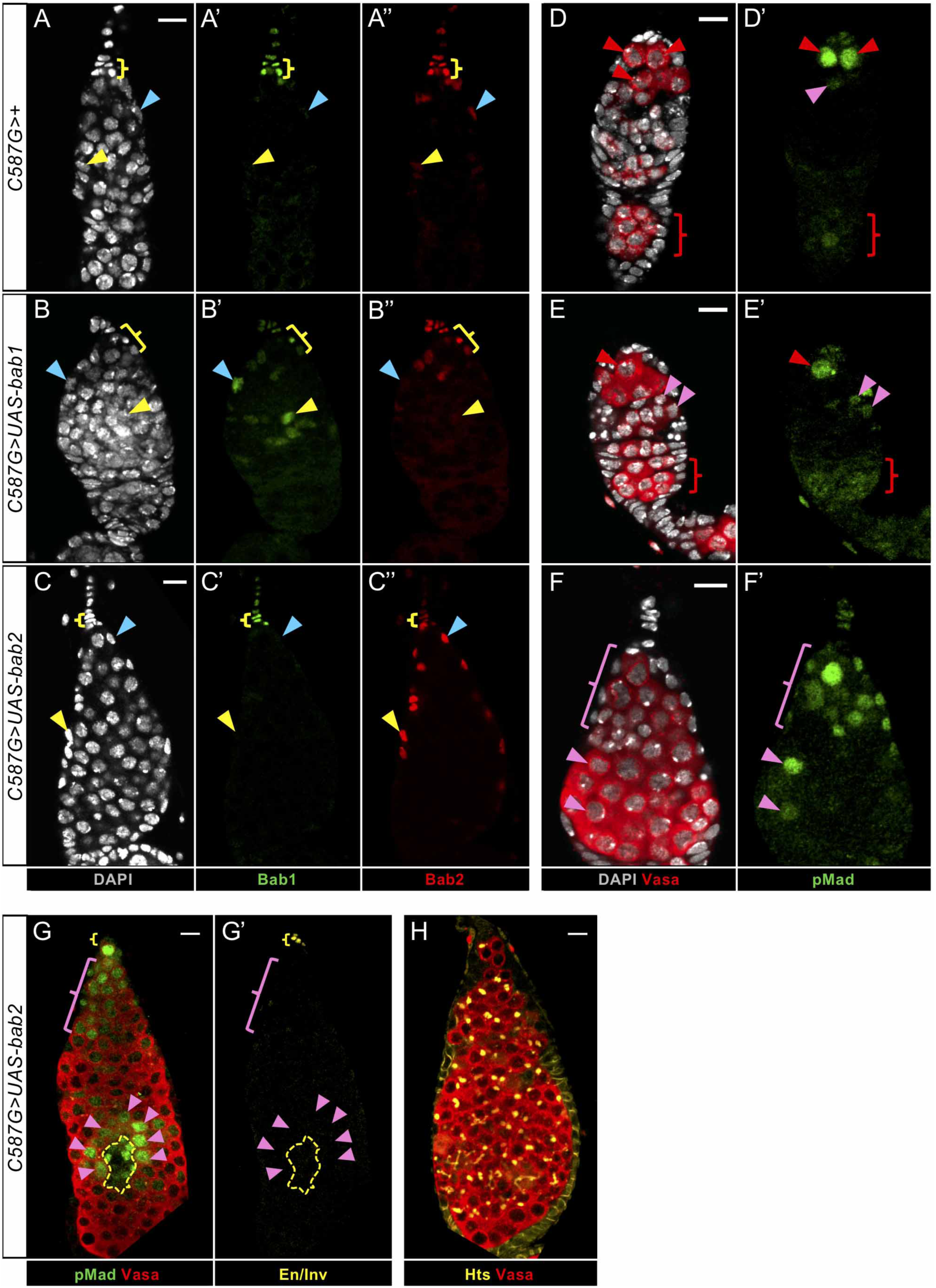
The overexpression of *bab2* is sufficient to induce GSC expansion in adult germaria. (A-H) Whole mount immunostaining of prepupal ovaries germaria from 10-day old females (projections of adjacent confocal sections). DAPI nuclear labeling Anterior is up. Scale bars: 10µm. (A-A”, D-D’) In a control (*C587G>+*), Bab1 (green, A’) and Bab2 (red, A’’) are present in niche cells, mainly in the Cap Cells (CCs) (yellow bracket) and overlying Terminal Filaments (TFs). Bab2 can also be detected faintly in some Escort Cells (ECs) (blue arrowhead, A’’) and more posterior somatic cells (yellow arrowhead, A”). (D,D’) Three Germinal Stem Cells (GSCs) are marked with pMad (green) at high or low level (red and pink arrowhead, respectively). pMad also accumulates faintly in the maturating germline cyst (red bracket). (B-F’) Ectopic expression of *bab1* (*C587G>UAS-bab1* - B’, green) or overexpression of *bab2* (*C587G>UAS-*bab1 - C”, red) causes elevated accumulation of the corresponding proteins in ECs and prefollicular cells compared to the control (B’,C” compare to A’,A”, blue and yellow arrowheads, respectively). Cross regulation between *bab1* and *bab2* does not occur (B’’ and C’, blue and yellow arrowheads). (E,E’) In rare cases, the ectopic expression of *bab1* leads to the presence of ectopic pMad+ Germ Cells (GCs) (pink arrowhead). pMad is also detected faintly in the maturating germline cyst as in the control (red bracket). (F-G) Inducing higher levels of Bab2 in ECs and prefollicular cells leads to an increase in the number of pMad+ GSCs in continuity with the niche (pink bracket) or at ectopic locations in germaria (pink arrowheads). (G,G’) Engrailed/ Invected are not detected in somatic cells (yellow dotted lines) in contact with the ectopic GSCs (pink arrowheads) compared to accumulation of these proteins in the endogenous niche (yellow bracket). (H) The overexpression of *bab2* leads to the formation of huge germaria with GSC-like tumors marked by the presence of spectrosome-containing (Hts, yellow) GCs (Vasa, red).

However, with respect to GCs (Vasa+), in ovaries overexpressing *bab2*, all germaria contained more than the normal number of GSCs as marked by the presence of pMad, some in continuity with the niche and others at ectopic locations (Fig 10F and 10F’ and Fig 10G, pink brackets and pink arrowheads, respectively, compared to the control in Fig 10D and 10D’). Among these germaria, 75% exhibited more than 20 GSCs and 25% between 5 and 20 GSCs (n=137), compared to 0 and 10 % (n=87), respectively, for control germaria. In many cases, overexpression of *bab2* led to the formation of huge germaria with GSC-like tumors as marked by the presence of spectrosomes (Hts+ cytoplasmic structures, Fig 10H). In these tumorous germaria, no ectopic En/Inv was detected in somatic cells outside of the endogenous niche (Fig 10G’, yellow dotted line and yellow bracket, respectively). Upon ectopic expression of *bab1*, only 4 germaria out of 133 exhibited ectopic GCs in which the BMP pathway was activated (Fig 10E and 10E’, pink arrowheads). Altogether, these results show that *bab2* overexpression in somatic cells outside of the adult GSC niche is sufficient to induce activation of the BMP pathway in adjacent GCs, thereby producing numerous ectopic GSCs and tumorous germaria. Since *bab1* ectopic expression did not affect GSC number significantly, either Bab1 does not have the equivalent capacity as Bab2, or is not expressed at the correct level in the condition tested to provoke an effect. In addition, since somatic Bab2 overexpression does not produce ectopic *En/Inv* expression, we conclude that Bab2 GSC-like tumor induction, associated with BMP pathway activation in GCs, does not require *en/inv* function in *bab2* overexpressing cells.

## DISCUSSION

### Bab 1 and Bab2 are redundant in precursor niche cells for functional GSC niche formation

The two *bab* paralogs encode transcription factors that have been shown to be required for leg proximal-distal differentiation, ovary morphogenesis and sexually-dimorphic abdomen pigmentation (Bartoletti et al., 2012; Cabrera et al., 2002; Godt and Laski, 1995; Kopp et al., 2000; Sahut-Barnola et al., 1995). *bab1* and *bab2* display partially overlapping expression patterns in these three organs (leg, ovary, abdomen) (Couderc et al., 2002; Kopp et al., 2000; Williams et al., 2008). Indeed, Bab1 is always present with Bab2, while Bab2 is present alone in additional territories in the leg and ovary (Couderc et al., 2002). The question of redundancy in function between Bab paralogs has been hampered in part by their overlapping expression (at least during leg and ovary development), but especially by the lack of any known *bab2* molecularly null mutation not affecting *bab1*. Indeed, for *bab1*, the *bab^A128^* allele has been shown to be associated with complete loss of Bab1 without affecting Bab2 levels (confirmed in the present study), but no equivalent allele has been characterized for *bab2* (Chalvet et al., 2011; Couderc et al., 2002). Two studies have now addressed the question of redundancy between the two *bab* genes using tissue-specific expression of *bab1* and *bab2* RNAi constructs, for abdomen pigmentation (Roeske et al., 2018) and the present study for ovarian GSC niche formation. Roeske *et al*. (Roeske et al., 2018) showed that both *bab1* and *bab2* are necessary for repression of pigmentation of female abdomen segments A5 and A6. However, since depletion of both proteins leads to a much stronger female pigmentation defect than depletion of each one separately, this and other results led the authors to propose that 4 doses of Bab proteins, provided in the wild-type by the presence of the two paralogs, are needed to repress expression of a pigmentation gene, *yellow*, in the female abdomen. In addition, their work and that of Couderc *et al*. (Couderc et al., 2002) indicate equivalent capacities for Bab1 and Bab2 since ectopic expression of either one at the same level in males leads to similar abnormal pigmentation of A5 and A6. Taken together, these results suggest redundancy between Bab1 and Bab2 for sexually-dimorphic abdominal pigmentation.

In developing ovary niches, our results indicate full redundancy between *bab1* and *bab2* for TF formation and strongly suggest that this is also the case for GSC establishment. Therefore, in ovarian GSC niches, two doses of Bab proteins seem sufficient for these processes. In contrast, ectopic and overexpression of *bab1* and *bab2,* respectively, outside of the GSC niche, did not produce similar effects. Indeed, somatic overexpression of *bab2* in germaria led to production of GSC-like tumors, while under similar conditions *bab1* did not significantly affect germaria. This may not be so surprising since in other contexts Bab proetin activities do not seem fully equivalent. For example, *bab* mutant rescue experiments conducted with respect to leg and ovary developmental mutant phenotypes have shown that both Bab1 and Bab2 are able to rescue the mutant phenotypes, but that Bab2 does so more efficiently (Bardot et al., 2002). In addition, *bab2* has a much larger expression domain than *bab1* in the ovary, and removing *bab2* from both GSC niches and the rest of the somatic cells has a much stronger effect on ovary morphogenesis than removing *bab2* only from niche cells (Green and Extavour, 2012). Altogether, *bab1* and *bab2* thus seem to have redundant functions in the tissues where they are co-expressed and *bab2* has additional functions in tissues where *bab1* is not expressed.

### A newly-identified role for Bab proteins in GSC establishment through *dpp* expression in CCs

The BMP family member Dpp, known to be emitted principally by CCs within niches germaria, has been shown to be essential for maintenance of GSC status in the adult (Luo et al., 2017; Song et al., 2004; Xie and Spradling, 1998). During larval to pupal transition, Dpp produced by niches has also been shown to be required for GSC establishment and proliferation (Gilboa and Lehmann, 2004; Matsuoka et al., 2013; Zhu and Xie, 2003). When both *bab1* and *bab2* functions were removed from precursor niche cells, these cells were not organized into TFs at the pupal stage and GSC establishment, as evidenced by activated Dpp signal transduction, was largely absent. Absence of GSCs does not seem to be due to a defect in CC specification, since the expression pattern of several markers, the combination of which distinguishes CCs, allowed us to identify cells presenting CC characteristics among *bab* deficient niche cells correctly positioned adjacent to GCs. Indeed, like wild-type CCs in prepupal ovaries, these cells produce Tj and En/Inv, express *P1444Z-lacZ* (albeit at somewhat lower levels than in control cells), are devoid of Delta at the plasma membrane but present rare Delta-containing vesicles and are activated for the Notch signaling pathway reporter *E(Spl)mβ-CD2.* Importantly, Tj and the Notch pathway have been shown to be necessary for CC specification (Hsu and Drummond-Barbosa, 2011; Panchal et al., 2017; Song et al., 2007; Yatsenko and Shcherbata, 2018). Therefore, several aspects of CC specification do not depend on *bab* gene functions.

However, in *bab* deficient niche cells expressing CC markers, the expression of two *dpp* transcriptional reporters, normally observed in control CCs, was absent. Therefore, Bab proteins act as activators of *dpp* transcription in CCs, directly or indirectly. Bab proteins are thus necessary for at least one very important property of CCs, that of expressing high levels of *dpp*, which is essential for their function as GSC niche cells (Liu et al., 2010, 2015; Luo et al., 2017; Wang et al., 2008; Xie and Spradling, 1998). Hence, *bab* requirement in GSC establishment likely relies on its control of *dpp* expression. Thus, our results reveal an as yet unknown function for Bab proteins in niches for GSC establishment.

TF cell specification seems more perturbed than that of CCs in Bab1 and Bab2-depleted niche cells. Though Tj expression was absent as in control TF cells, En/Inv levels were two-fold lower, and *P1444-lacZ* expression and membranous Delta were undetectable, unlike in control TF cells. It is therefore not possible to attribute a clear identity to these cells. Incorrect TF cell specification could explain why these cells are unable to undergo proper TF morphogenesis. Taken together these results indicate that in developing niches Bab proteins are necessary for TF cell specification and for the functionality of CCs for GSC establishment.

### En/Inv are not essential for TF formation nor GSC establishment, but seem necessary for GSC maintenance as of niche formation

In the ovary, *en/inv* are expressed specifically in niches during their formation. Our results show that their functions, however, are not necessary for niche formation. Indeed, we found that when En/Inv were depleted in developing niches by RNAi, the same number of correctly formed TFs, TF cells and CCs were found as in controls. This is consistent with the observation of normal larval ovaries upon En/Inv depletion in Allbee *et al*. (Allbee et al., 2018), using a different niche cell Gal4 driver and the same *en/inv* RNAi transgenes as used in our study. In contrast, another study, based on induction of clones of ovarian somatic cells homozygous for a null allele for both *en/inv*, concluded that these genes are involved specifically for proper TF cell alignment within individual stacks in larval ovaries (Bolívar et al., 2006). The difference with the results presented here, may be related to the different approaches used to abolish *en/inv* expression. With the RNAi approach used here, all niche cells are depleted of En/Inv, while *en/inv* null mutant clone induction used in Bolívar *et al*. (Bolívar et al., 2006) led to the formation of stacks with both wild-type and mutant TF cells. Possible heterogeneity in the identity between the two populations of cells may have impeded normal interaction between the cells during the stacking of TF cells.

Our results also indicate that En/Inv are not essential for initial GSC establishment since when they were strongly reduced, the proportion of prepupal niches associated with at least one GSC was the same as in the control. However, if left to develop until eclosion, these ovaries had lost all their GSCs suggesting that GSCs could not be maintained without the presence of En/Inv in niche cells. When expression of *en/inv* was restored in niche cells from the beginning of pupariation, then GSCs were present in adult germaria. These results suggest that the known function of En/Inv for GSC maintenance in the adult (Luo et al., 2017; Rojas-Ríos et al., 2012), probably through direct regulation of *dpp* expression (Luo et al., 2017), may be additionally necessary for this process earlier, as of niche formation during larval/prepupal stages. Indeed, at the prepupal stages the overall number of GSCs upon En/Inv depletion was already about 30% less than that in the control, and these GSCs showed a globally lower level of BMP signal activation than that in the control. We propose that Bab proteins are essential for initial GSC establishment through control of *dpp* expression in CCs during larval stages, and that as of this point, En/Inv are required for GSC maintenance, likely also through *dpp* regulation in these same cells. Therefore, taken together, our results suggest that the molecular players responsible for regulation of *dpp* expression in CCs as of niche formation, first for GSC establishment and, subsequently, for GSC maintenance may not be the same. Consistent with this, the absence of Bab proteins in CCs does not affect En/Inv levels in these same cells, and GSC establishment is nonetheless highly affected. However, it is not possible to completely exclude that En/Inv participate to *dpp* expression and GSC establishment and/or proliferation, in parallel to Bab, but to a lesser extent than Bab. Indeed, in prepupal ovaries, the region depleted of both Bab proteins is associated with the presence of almost no GSCs, while upon En/Inv depletion at this same stage, about two thirds of the normal number of GSCs are present.

A newly-identified gene necessary for ovary morphogenesis is *Lmx1a*, encoding a LIM-homeodomain transcription factor expressed in somatic apical cells, TF cells and CCs in the prepupal ovary (Allbee et al., 2018). Transcriptomic analysis revealed that *en/inv* are positively regulated by *Lmx1a* in ovaries at the larval to pupal transition, but Lmx1a was shown to be necessary for proper organization and differentiation of niche cells, unlike En/Inv. Indeed, *Lmx1a* niche mutant phenotypes resemble those of *bab* genes and strong genetic interactions were demonstrated between *Lmx1a* and *bab* genes. In addition, Bab proteins were shown to be necessary for *Lmx1a* expression and thus Lmx1a is a potential Bab target, direct or indirect, to be tested in the future for mediation of the role of Bab in niches for their correct formation.

### *bab2* overexpression leads to production of GSC-like ovarian tumors in adults

We have shown that in developing niches Bab proteins are necessary for BMP pathway activation and GSC establishment. We therefore also addressed whether ectopic Bab proteins in somatic cells outside of the niche could be sufficient to induce BMP pathway activation in adjacent GCs. During ovary development, we did not detect a significant effect upon *bab1* ectopic expression or *bab2* overexpression in non-niche somatic cells. In contrast, in adult germaria, overexpression of *bab2* in ECs and prefollicular cells led to the formation of GSC-like germarial tumors indicating that Bab2 is sufficient for acquisition of GSC status. The GSC-like germarial tumor phenotype was similar to that previously described for germarial overexpression of *dpp* (Song et al., 2004; Xie and Spradling, 1998) or *en* (Eliazer et al., 2014; Luo et al., 2017). Importantly, the phenotype observed in germaria overexpressing *bab2* was not a consequence of ectopic expression of En/Inv, since in these germaria, En/Inv were found only in endogenous niches as in the control, far from ectopic GSCs. Thus, ectopic Dpp activity in GCs induced by *bab2* overexpression is independent of *en/inv* function. Therefore, our results indicate that Bab controls Dpp pathway activation both in GSCs of endogenous niches and in ectopic GSCs in adult germaria, and that, in both cases, this is independent of En/Inv. Since we show here that *bab* is necessary for *dpp* expression in CCs and for GSC recruitment during niche formation, it is possible that the presence of ectopic GSCs in adult germaria overexpressing *bab2* somatically could be due to ectopic activation of *dpp* expression in these cells. Interestingly, inhibition of the JAK/STAT pathway in niche cells has also been shown to result in reduction in GSC numbers associated with low *dpp* expression in niches, without affecting expression of other CC markers (Decotto and Spradling, 2005; Lopez-Onieva et al., 2008; Wang et al., 2008). In addition, ectopic activation of the JAK/STAT pathway in the germarium is sufficient to induce ectopic GSC-like cells associated with ectopic *dpp* expression but not with ectopic CC specification. Bab interaction with this signaling pathway remains to be explored.

## MATERIAL AND METHODS

### Fly stocks

We used *hedgehog-Gal4* (gift from P. Therond and hereafter called *hhG*), a Gal4-expressing enhancer trap insertion in the *hh* locus, to target niche cells during their differentiation, and *bab^PGal4-2^* (gift from J.L. Couderc and hereafter called *babG*), a Gal4-expressing enhancer trap insertion in the *bab l*ocus, to drive expression in all somatic cells of the larval ovary (Cabrera et al., 2002) and present study). These drivers were combined with *UAS-dicer2* and *tub-Gal80^TS^* (gift from J. Montagne) when indicated. Two different *UAS-GFP* transgenes (one associated with a Nuclear Localization Signal sequence, Bloomington Drosophila Stock Center, BDSC 4476, and one without, gift from A. Boivin), were used in combination with the *hhG* driver to mark niche cells. *The* RNAi lines *UAS-bab1^IR^* (Vienna Drosophila Stock Center, VDRC 50285) and *UAS-bab2^IR^* (VDRC 49042), and the *bab1* null allele *bab^A128^* (Godt et al., 1993) (gift from D.Godt) were used for the *bab* loss-of-function analysis. *bab^AR07^,FRT80B/TM6B* (gift from M. Boube) and *hs-FLP;; ubi-GFP, FRT80B* (gift from J. Montagne) lines were used to generate *bab* null mitotic cell clones. *bab^AR07^* is a deletion inactivating both *bab1* and *bab2* (Couderc et al., 2002). *UAS-en^IR^* (VDRC 35697) and *UAS-inv^IR^* (BDSC 41675) were used for the *engrailed/invected* knockdown analysis. The enhancer reporter line *P{PZ}1444* (*P1444-lacZ*) (Margolis and Spradling, 1995) and *bamP-GFP*, a GFP transcriptional reporter for *bam* expression ((Chen and McKearin, 2003a), Kyoto DGRC 118177) were used as cell-type specific markers. *E(spl)mβ-CD2*, in which the sequence encoding rat CD2 protein is inserted downstream of the *Enhancer of Split* [*E(spl)mβ*] promoter that is activated by Notch signaling, was used as a readout of Notch pathway activity (de Celis et al., 1998) (gift from A. Bardin). *dpp-GFP* (VDRC 318464) and *dpp-nlsGFP* (VDRC 318414) (Sarov et al., 2016) and *P4-lacZ* (Li et al., 2016) (gift from R. Xi) were used as *dpp* transcriptional reporters. *dpp-nlsGFP* contains 43766bp of the *dpp* genomic locus. *P4-lacZ* contains 1494bp of a *dpp* enhancer region. To drive ectopic and over-expression of *bab1* and *bab2*, we used *C587-Gal4* (Zhu and Xie, 2003) (gift from T. Xie), *UAS-bab1* (BDSC, 6939) and *UAS-bab2^4-66^* (Bardot et al., 2002; Couderc et al., 2002) (gift from A. Kopp).

### Experimental conditions

For all the experiments, flies were raised on standard cornmeal food.

For pupal analysis of the effect of *bab* and *engrailed/invected* (*en/inv*) depletion in niche cells with *bab^A128^*, *hhG*, *babG*, and RNAi lines, crosses were started at 25°C, parents removed 24-to-48h later, descendants were then transferred to 29°C and ovaries from 1-to-6 hour old prepupae (white pupae) were dissected.

To generate *bab* mosaic mutant prepupal ovaries, *bab^AR07^, FRT80B/TM6B* flies were crossed to *hs-FLP;; ubi-GFP, FRT80B* flies at 25°C. Clones marked by absence of GFP were induced by one 1-hour heat shock at 38°C at the end of the second instar larval stage, and prepupae were dissected for ovary analysis 50h after heat shock.

Transient expression of the *UAS-engrailed^IR^* and *UAS-invected^IR^* transgenes was achieved using *Tub Gal80^TS^ and babG.* For analysis of adult (1-day old) ovaries upon RNAi-mediated knockdown of *engrailed/invected* (*en/inv*), two conditions were used: 18°C (from crosses to dissection) for the control and 29°C (crosses at 25°C and transferred 24-to-48h later to 29°C) for the depletion of En/Inv. To test the effect of the depletion only during niche formation, *en/inv* were reexpressed at the prepupal stage by shifting prepupae from 29°C to 18°C for the rest of development.

To analyze the effect, at prepupal stage, of *bab1* ectopic expression and *bab2* overexpression in somatic cells of the ovaries during larval stages, crosses were conducted at 25°C for 24-to-48h and descendants were then transferred either to 29°C or maintained at 25°C until dissection.

For *bab1* and *bab2* gain-of-function experiments in the adult, individuals were raised at 18°C and then shifted to 29°C after eclosion. Ovaries from 10 day-old females were dissected.

### Immunostaining

Prepupal ovaries were dissected in PBS medium, fixed in 4% formaldehyde (R1026, Agar Scientific) for 20 to 30 minutes, and washed 3 x 10 minutes in PBT (PBS supplemented with 0.3% tritonX-100 – Sigma T8787). Ovaries were then incubated in a blocking solution (PBTA: PBT supplemented with 1% BSA – Sigma A3059) for a minimum of 20 minutes. Primary antibodies were diluted in PBTA and ovaries were incubated in this solution for 6 hours at room temperature or overnight at 4°C. The following primary antibodies were used: rabbit anti-Smad3 (1:200, ab52903, Abcam); rabbit anti-Bab1 (1:4000, gift from T. Williams); rabbit anti-GFP (1:200, FP-37151B, Interchim); rat anti-Bab2 (1:4000, gift from J-L. Couderc); rat IgM anti-Vasa (1:500, Developmental Studies Hybridoma Bank-DSHB); mouse anti-En/Inv recognizing both paralogs (1:200, 4D9, DSHB); guinea-pig anti GFP (1:200, 132 002, Synaptic System); mouse anti-β-Gal (1:400, 40-1a, DSHB,); guinea pig anti-Traffic Jam (1:5000, gift from D. Godt); mouse anti-Delta (1:200, C594.9B, DSHB), mouse anti-rat CD2 (1:100, MCA154GA, Biorad); mouse anti hts (1:200, DSHB). After primary antibody incubation, tissues were washed in PBT 3×10min, and incubated in PBTA for at least 30 minutes before incubation in secondary antibodies in PBTA. The following secondary antibodies were used: donkey anti-mouse Cy3 (715-165-151-Jackson Laboratories); and the following ones from Thermo Fisher Scientific: chicken anti-rabbit 488 (A21441); goat anti-rabbit 568 (A11011); goat anti-guinea pig 488 (A11073), anti-mouse 488 (A11029); anti-rat IgM 647 (A21248); anti-rat IgG 647 (A21247). Nuclei were detected with DAPI (1:200, 1mg/ml) and F-actin with Phalloidin:647 (1:200, 65906, Sigma). After secondary antibody incubation, tissues were washed 3×10 min in PBT, and mounted between slide and coverslip in DABCO (D27602, Sigma) with 70% glycerol. For larval ovaries, a spacer was positioned between the slide and the coverslip to avoid ovary squashing. The lateral side of the prepupal ovary was identified by its contact with the fat body.

### Fluorescence microscopy, image analysis and statistics

Images were acquired with a 40x or 63x objective on a confocal laser-scanning microscope Leica TCS SP8 equipped with 405nm, 488nm, 552nm, and 638nm emission diodes. Same settings were used between the different genotypes in each experiment. Prepupal ovaries were acquired with 1µm steps between each section and AOTF/EOF compensation was used to increase diode percentage during acquisition using the LasX software. Adult ovary images were acquired with 0.5 or 1µm steps between each section.

Images represent either a projection of 4 consecutive sections or a 3D reconstruction of the entire tissue. Images were processed with Fiji (Schindelin et al., 2012).

Counting TF number in prepupal ovaries was achieved through immunodetection of niche cell markers, such as GFP (*hhG+* cells), Bab1 or En/Inv, or DAPI labeling of flattened TF cell nuclei. Both lateral views and 3D reconstructions from the anterior pole of ovaries were used for this quantification. To distinguish TF cells and CCs among niche cells, labeling of F-actin which delimits cell contours and/or DAPI labeling of flattened TF cell nuclei was used. Niche cells with contours covering the entire width of the TF and with flattened nuclei were considered as TF cells, and more basal cells contacting GCs that did not cover the width of the TF and had round nuclei, as CCs (See Fig1B, compare green and yellow brackets, respectively).

*hhG+* cell flattening of control prepupal ovaries (*hhG>GFP*) or upon depletion of Bab proteins (*hhG>GFP, bab1^IR^, bab2^IR^*) was quantified with GFP immunostaining marking *hhG+* cells and fluorescent phalloïdin-bound F-actin marking cell contours in projections of 5 consecutive sections separated by 1µm. The width and height between cell contours of individual *hhG+* cells were measured independently, and the degree of flattening corresponded to the ratio between the width and height.

For comparison of En/Inv levels between control niche cells and niche cells depleted of Bab proteins (*bab^AR07^* null allele and RNAi-mediated depletion), the confocal section presenting the biggest area for individual nuclei was used to quantify fluorescence intensity of En/Inv levels using the RawIntDen tool in Fiji.

To quantify GSC number in both prepupal ovaries and adult niches, we used nuclear pMad staining in GCs as a readout of Dpp pathway activity. pMad staining levels in nuclei were very variable between GCs of same ovary and between ovaries of a same experiment ranging from very high to very low when detected with Royal LUT. GCs presenting low but unambiguous nuclear pMad staining were considered separately from GCs with high pMad+ GCs. GCs with high or low pMad levels were both considered as GSCs since pMad detection reflects a high or moderate activity of Dpp/BMP pathway (see Figure 1C-C’’).

Prism 7 (GraphPad Software) was used for statistical tests and for the generation of graphs. Values are presented as means + s.e.m., p-value calculated using a two-tailed Student’s t-test when two conditions were compared or an ANOVA one-way test for comparisons between multiple conditions. Results are indicated with: NS (Not Significant - p>0.05); * (0.05>p>0.01); ** (0.01>p>0.001); *** (0.001>p>0.0001) and **** (p<0.0001).

---

S1 Fig. **Expression pattern of *dpp-GFP* and *dpp-nlsGFP* transcriptional reporters in adult germaria.** Whole mount immunostaining of adult germaria (projections of adjacent confocal sections) from females raised at 25°C for detection of GFP (green), Engrailed/Invected (En/Inv, red) and Traffic Jam/Vasa (Tj/Vasa, magenta). Nuclei are labeled with DAPI (grey). Anterior is up. Scale bars: 10µm. (A,B) Germaria expressing *dpp-GFP* and *dpp-nlsGFP*, respectively. (A’-A’’’, B’-B’’’) Higher magnification of the regions framed with dotted lines in the corresponding germaria. For both constructions, GFP (green) is present in the Cap Cells (yellow brackets) co-marked by nuclear Engrailed/Invected (En/Inv, red) and nuclear Traffic Jam (TJ, magenta), and in some Escort Cells (blue brackets), but absent from Terminal Filament cells (green brackets) marked by En/Inv (red). Expression of both GFP reporters was also detected in prefollicular cells (orange brackets).

S2 Fig. **The ectopic expression of *bab1* or the overexpression of *bab2* does not affect ovary morphogenesis.** (A-C) Whole mount immunostaining of prepupal ovaries (projections of adjacent confocal sections) from females raised at 25°C. Anterior is up and medial is left. Scale bars: 10µm. Nuclei are labeled with DAPI (grey). (A-A’’) Control ovary (*C587-Gal4* (*C587G)>+*) showing the expression pattern of *bab1* (strong expression in niche cells and weak in Intermingled Cells (ICs, white brackets). (B-B’’) The *C587G* driver coupled with *UAS-bab1* allows the ectopic expression of *bab1* in some ICs (arrowheads). (C-C’’) The *C587G* driver coupled with *UAS-bab2* allows an overexpression of *bab2* in some ICs (arrowheads). (B’’,C’’) Neither of these constructs causes ectopic accumulation of En/Inv (red) in ICs overexpressing *bab1* or *bab2* (arrowheads). The overall organization of these ovaries does not seem disturbed compared to the control.

## REFERENCES

Aguado, B.A., Bushnell, G.G., Rao, S.S., Jeruss, J.S., and Shea, L.D. (2017). Engineering the pre-metastatic niche. Nat. Biomed. Eng. 1.

Allbee, A.W., Rincon-Limas, D.E., and Biteau, B. (2018). Lmx1a is required for the development of the ovarian stem cell niche in *Drosophila*. Development 145, dev163394.

Bardot, O., Godt, D., Laski, F.A., and Couderc, J.-L. (2002). Expressing UAS-bab1 and UAS-bab2: A comparative study of gain-of-function effects and the potential to rescue the bric à brac mutant phenotype. Genesis 34, 66–70.

Bartoletti, M., Rubin, T., Chalvet, F., Netter, S., Dos Santos, N., Poisot, E., Paces-Fessy, M., Cumenal, D., Peronnet, F., Pret, A.-M., et al. (2012). Genetic Basis for Developmental Homeostasis of Germline Stem Cell Niche Number: A Network of Tramtrack-Group Nuclear BTB Factors. PLoS ONE 7.

Bolívar, J., Pearson, J., López-Onieva, L., and González-Reyes, A. (2006). Genetic dissection of a stem cell niche: The case of the Drosophila ovary. Dev. Dyn. 235, 2969–2979.

Cabrera, G.R., Godt, D., Fang, P.-Y., Couderc, J.-L., and Laski, F.A. (2002). Expression pattern of Gal4 enhancer trap insertions into the bric à brac locus generated by P element replacement. Genesis 34, 62–65.

de Celis, J.F., Tyler, D.M., de Celis, J., and Bray, S.J. (1998). Notch signalling mediates segmentation of the Drosophila leg. 10.

Chaharbakhshi, E., and Jemc, J.C. (2016). Broad-complex, tramtrack, and bric-à-brac (BTB) proteins: Critical regulators of development. Genesis 54, 505–518.

Chalvet, F., Bartoletti, M., and Théodore, L. (2011). Ovary phenotype and expression of bab1 and bab2paralogs in the ovary of two mutants of bab locus in Drosophila melanogaster. Drosoph Info Serv 158–162.

Chen, D., and McKearin (2003a). A discrete transcriptional silencer in the bam gene determines asymmetric division of the Drosophila germline stem cell. Development 130, 1159–1170.

Chen, D., and McKearin, D. (2003b). Dpp Signaling Silences bam Transcription Directly to Establish Asymmetric Divisions of Germline Stem Cells. Curr. Biol. 13, 1786–1791.

Cohen, E.D., Mariol, M.-C., Wallace, R.M.H., Weyers, J., Kamberov, Y.G., Pradel, J., and Wilder, E.L. (2002). DWnt4 Regulates Cell Movement and Focal Adhesion Kinase during Drosophila Ovarian Morphogenesis. Dev. Cell 2, 437–448.

Couderc, J.-L., Godt, D., Zollman, S., Chen, J., Li, M., Tiong, S., Cramton, S.E., Sahut-Barnola, I., and Laski, F.A. (2002). The bric à brac locus. 15.

Decotto, E., and Spradling, A.C. (2005). The Drosophila ovarian and testis stem cell niches: similar somatic stem cells and signals. Dev. Cell 9, 501–510.

Eliazer, S., Palacios, V., Wang, Z., Kollipara, R.K., Kittler, R., and Buszczak, M. (2014). Lsd1 restricts the number of germline stem cells by regulating multiple targets in escort cells. PLoS Genet. 10, e1004200.

Ermolaeva, M., Neri, F., Ori, A., and Rudolph, K.L. (2018). Cellular and epigenetic drivers of stem cell ageing. Nat. Rev. Mol. Cell Biol. 19, 594.

Forbes, A.J., Spradling, A.C., Ingham, P.W., and Lin, H. (1996). The role of segment polarity genes during early oogenesis in Drosophila. 12.

Gancz, D., Lengil, T., and Gilboa, L. (2011). Coordinated Regulation of Niche and Stem Cell Precursors by Hormonal Signaling. PLoS Biol. 9.

Gilboa, L. (2015). Organizing stem cell units in the Drosophila ovary. Curr. Opin. Genet. Dev. 32, 31–36.

Gilboa, L., and Lehmann, R. (2004). Repression of Primordial Germ Cell Differentiation Parallels Germ Line Stem Cell Maintenance. Curr. Biol. 14, 981–986.

Godt, D., and Laski, F.A. (1995). Mechanisms of cell rearrangement and cell recruitment in Drosophila ovary morphogenesis and the requirement of bric à brac. 15.

Godt, D., Couderc, J.-L., Cramton, S.E., and Laski, F.A. (1993). Pattern formation in the limbs of Drosophila: bric à brac is expressed in both a gradient and a wave-like pattern and is required for specification and proper segmentation of the tarsus. 14.

Green, D.A., and Extavour, C.G. (2012). Convergent evolution of a reproductive trait through distinct developmental mechanisms in Drosophila. Dev. Biol. 372, 120–130.

Green, D.A., Sarikaya, D.P., and Extavour, C.G. (2011). Counting in oogenesis. Cell Tissue Res. 344, 207–212.

Greenspan, L.J., de Cuevas, M., and Matunis, E. (2015). Genetics of Gonadal Stem Cell Renewal. Annu. Rev. Cell Dev. Biol. 31, 291–315.

Gustavson, E., Goldsborough, A.S., Ali, Z., and Kornberg, T.B. (1996). The Drosophila engrailed and invected genes: partners in regulation, expression and function. Genetics 142, 893–906.

Hsu, H.-J., and Drummond-Barbosa, D. (2011). Insulin signals control the competence of the Drosophila female germline stem cell niche to respond to Notch ligands. Dev. Biol. 350, 290–300.

Kai, T., and Spradling, A. (2004). Differentiating germ cells can revert into functional stem cells in Drosophila melanogaster ovaries. Nature 428, 564–569.

Kaplan, R.N., Riba, R.D., Zacharoulis, S., Bramley, A.H., Vincent, L., Costa, C., MacDonald, D.D., Jin, D.K., Shido, K., Kerns, S.A., et al. (2005). VEGFR1-positive haematopoietic bone marrow progenitors initiate the pre-metastatic niche. Nature 438, 820–827.

Kopp, A., Duncan, I., and Carroll, S.B. (2000). Genetic control and evolution of sexually dimorphic characters in Drosophila. Nature 408, 553–559.

Lai, C.-M., Lin, K.-Y., Kao, S.-H., Chen, Y.-N., Huang, F., and Hsu, H.-J. (2017). Hedgehog signaling establishes precursors for germline stem cell niches by regulating cell adhesion. J. Cell Biol. 216, 1439–1453.

Li, M.A., Alls, J.D., Avancini, R.M., Koo, K., and Godt, D. (2003). The large Maf factor Traffic Jam controls gonad morphogenesis in Drosophila. Nat. Cell Biol. 5, 994–1000.

Li, X., Yang, F., Chen, H., Deng, B., Li, X., and Xi, R. (2016). Control of germline stem cell differentiation by Polycomb and Trithorax group genes in the niche microenvironment. Dev. Camb. Engl. 143, 3449–3458.

Liu, M., Lim, T.M., and Cai, Y. (2010). The Drosophila Female Germline Stem Cell Lineage Acts to Spatially Restrict DPP Function Within the Niche. Sci. Signal. 3, ra57–ra57.

Liu, Z., Zhong, G., Chai, P.C., Luo, L., Liu, S., Yang, Y., Baeg, G.-H., and Cai, Y. (2015). Coordinated niche-associated signals promote germline homeostasis in the Drosophila ovary. J. Cell Biol. 211, 469–484.

Lopez-Onieva, L., Fernandez-Minan, A., and Gonzalez-Reyes, A. (2008). Jak/Stat signalling in niche support cells regulates dpp transcription to control germline stem cell maintenance in the Drosophila ovary. Development 135, 533–540.

Lu, T., Wang, S., Gao, Y., Mao, Y., Yang, Z., Liu, L., Song, X., Ni, J., and Xie, T. (2015). COP9-Hedgehog axis regulates the function of the germline stem cell progeny differentiation niche in the Drosophila ovary. Development 142, 4242–4252.

Luo, L., Siah, C.K., and Cai, Y. (2017). Engrailed acts with Nejire to control *decapentaplegic* expression in the *Drosophila* ovarian stem cell niche. Development 144, 3224–3231.

Margolis, J., and Spradling, A. (1995). Identification and behavior of epithelial stem cells in the Drosophila ovary. Dev. Camb. Engl. 121, 3797–3807.

Matsuoka, S., Hiromi, Y., and Asaoka, M. (2013). Egfr signaling controls the size of the stem cell precursor pool in the Drosophila ovary. Mech. Dev. 130, 241–253.

McKearin, D., and Ohlstein, B. (1995). A role for the Drosophila bag-of-marbles protein in the differentiation of cystoblasts from germline stem cells. Development 121, 2937–2947.

Mendes, C.C., and Mirth, C.K. (2016). Stage-Specific Plasticity in Ovary Size Is Regulated by Insulin/Insulin-Like Growth Factor and Ecdysone Signaling in Drosophila. Genetics 202, 703–719.

Panchal, T., Chen, X., Alchits, E., Oh, Y., Poon, J., Kouptsova, J., Laski, F.A., and Godt, D. (2017). Specification and spatial arrangement of cells in the germline stem cell niche of the Drosophila ovary depend on the Maf transcription factor Traffic jam. PLoS Genet. 13.

Plaks, V., Kong, N., and Werb, Z. (2015). The Cancer Stem Cell Niche: How Essential Is the Niche in Regulating Stemness of Tumor Cells? Cell Stem Cell 16, 225–238.

Prager, B.C., Xie, Q., Bao, S., and Rich, J.N. (2019). Cancer Stem Cells: The Architects of the Tumor Ecosystem. Cell Stem Cell 24, 41–53.

Roeske, M.J., Camino, E.M., Grover, S., Rebeiz, M., and Williams, T.M. (2018). Cis-regulatory evolution integrated the Bric-à-brac transcription factors into a novel fruit fly gene regulatory network. ELife 7.

Rojas-Ríos, P., Guerrero, I., and González-Reyes, A. (2012). Cytoneme-Mediated Delivery of Hedgehog Regulates the Expression of Bone Morphogenetic Proteins to Maintain Germline Stem Cells in Drosophila. PLoS Biol. 10.

Sahut-Barnola, Dastugue, Bernard, and Couderc, J.-L. (1996). (PDF) Terminal filament cell organization in the larval ovary of Drosophila melanogaster: Ultrastructural observations and pattern of divisions.

Sahut-Barnola, I., Godt, D., Laski, F.A., and Couderc, J.-L. (1995). Drosophila Ovary Morphogenesis: Analysis of Terminal Filament Formation and Identification of a Gene Required for This Process. Dev. Biol. 170, 127–135.

Sarikaya, D.P., and Extavour, C.G. (2015). The Hippo Pathway Regulates Homeostatic Growth of Stem Cell Niche Precursors in the Drosophila Ovary. PLoS Genet. 11.

Sarikaya, D.P., Belay, A.A., Ahuja, A., Dorta, A., Green, D.A., and Extavour, C.G. (2012). The roles of cell size and cell number in determining ovariole number in Drosophila. Dev. Biol. 363, 279–289.

Sarov, M., Barz, C., Jambor, H., Hein, M.Y., Schmied, C., Suchold, D., Stender, B., Janosch, S., KJ, V.V., Krishnan, R., et al. (2016). A genome-wide resource for the analysis of protein localisation in Drosophila. ELife 5, e12068.

Sato, T., Ogata, J., and Niki, Y. (2010). BMP and Hh Signaling Affects Primordial Germ Cell Division in Drosophila. Zoolog. Sci. 27, 804–810.

Schindelin, J., Arganda-Carreras, I., Frise, E., Kaynig, V., Longair, M., Pietzsch, T., Preibisch, S., Rueden, C., Saalfeld, S., Schmid, B., et al. (2012). Fiji: an open-source platform for biological-image analysis. Nat. Methods 9, 676–682.

Song, X., zhu, and Xie (2002). Germline Stem Cells Anchored by Adherens Junctions in the Drosophila Ovary Niches. Science 296, 1855–1857.

Song, X., wong, and kawase (2004). Bmp signals from niche cells directly repress transcription of a differentiation-promoting gene, bag of marbles, in germline stem cells in the Drosophila ovary. Development 131, 1353–1364.

Song, X., Call, G.B., Kirilly, D., and Xie, T. (2007). Notch signaling controls germline stem cell niche formation in the Drosophila ovary. Development 134, 1071–1080.

Tseng, C.-Y., Su, Y.-H., Yang, S.-M., Lin, K.-Y., Lai, C.-M., Rastegari, E., Amartuvshin, O., Cho, Y., Cai, Y., and Hsu, H.-J. (2018). Smad-Independent BMP Signaling in Somatic Cells Limits the Size of the Germline Stem Cell Pool. Stem Cell Rep. 11, 811–827.

Wang, X., and Page-McCaw, A. (2018). Wnt6 maintains anterior escort cells as an integral component of the germline stem cell niche. Dev. Camb. Engl. 145.

Wang, L., Li, Z., and Cai, Y. (2008). The JAK/STAT pathway positively regulates DPP signaling in the Drosophila germline stem cell niche. J. Cell Biol. 180, 721–728.

Ward, E.J., Shcherbata, H.R., Reynolds, S.H., Fischer, K.A., Hatfield, S.D., and Ruohola-Baker, H. (2006). Stem Cells Signal to the Niche through the Notch Pathway in the Drosophila Ovary. Curr. Biol. 16, 2352–2358.

Williams, T.M., Selegue, J.E., Werner, T., Gompel, N., Kopp, A., and Carroll, S.B. (2008). The Regulation and Evolution of a Genetic Switch Controlling Sexually Dimorphic Traits in Drosophila. Cell 134, 610–623.

Xie, T., and Spradling, A.C. (1998). decapentaplegic Is Essential for the Maintenance and Division of Germline Stem Cells in the Drosophila Ovary. Cell 94, 251–260.

Xie, T., and Spradling, A.C. (2000). A Niche Maintaining Germ Line Stem Cells in the Drosophila Ovary. Science 290, 328–330.

Yatsenko, A.S., and Shcherbata, H.R. (2018). Stereotypical architecture of the stem cell niche is spatiotemporally established by miR-125-dependent coordination of Notch and steroid signaling. Development 145, dev159178.

Zhao, dong, and li (2018). Targeting cancer stem cells and their niche_ perspectives for future therapeutic targets and strategies | Elsevier Enhanced Reader.

Zhu, C.-H., and Xie, T. (2003). Clonal expansion of ovarian germline stem cells during niche formation in Drosophila. Development 130, 2579–2588.

