## Supplementary figures and images for "The Bric-à-Brac transcription factors are necessary for formation of functional germline stem cell niches through control of *dpp* expression in the *Drosophila melanogaster* ovary"

### S1 Fig

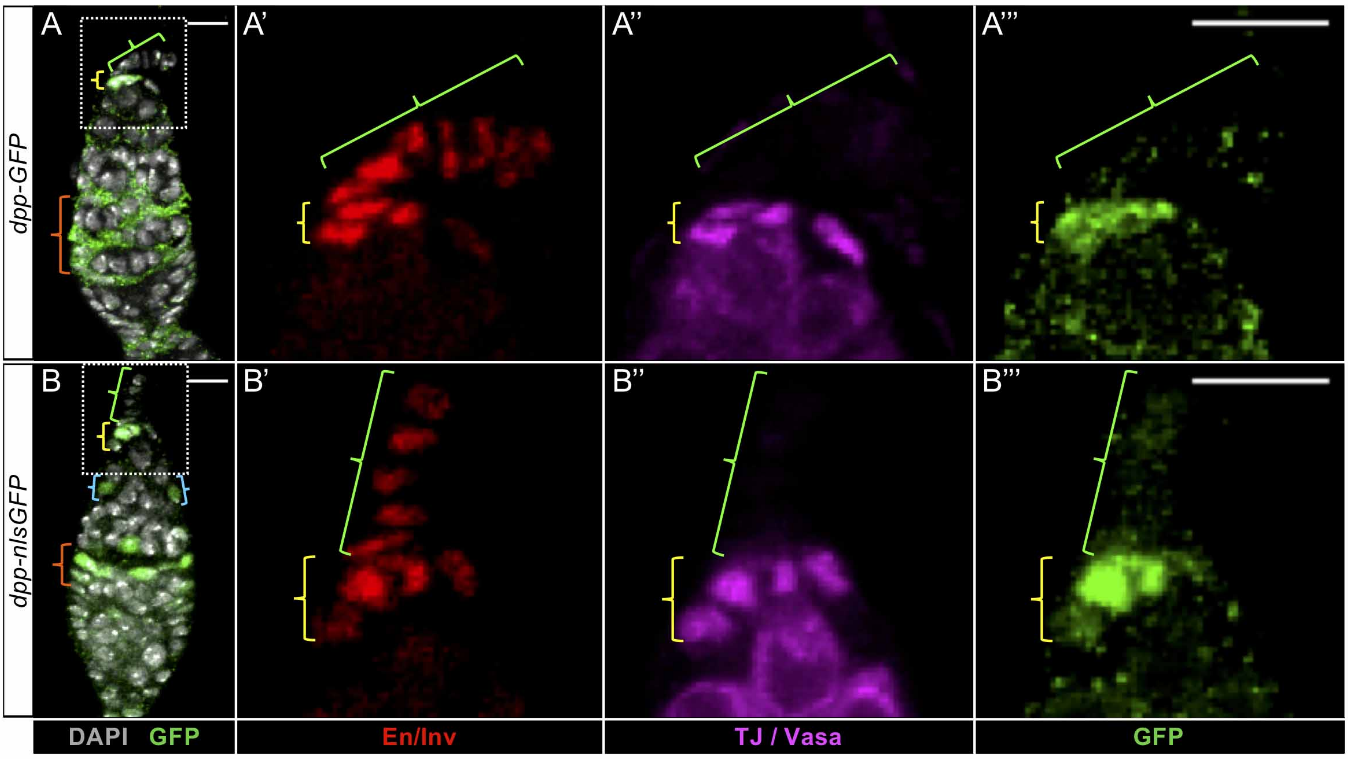


Supp Fig 1

### S2 Fig

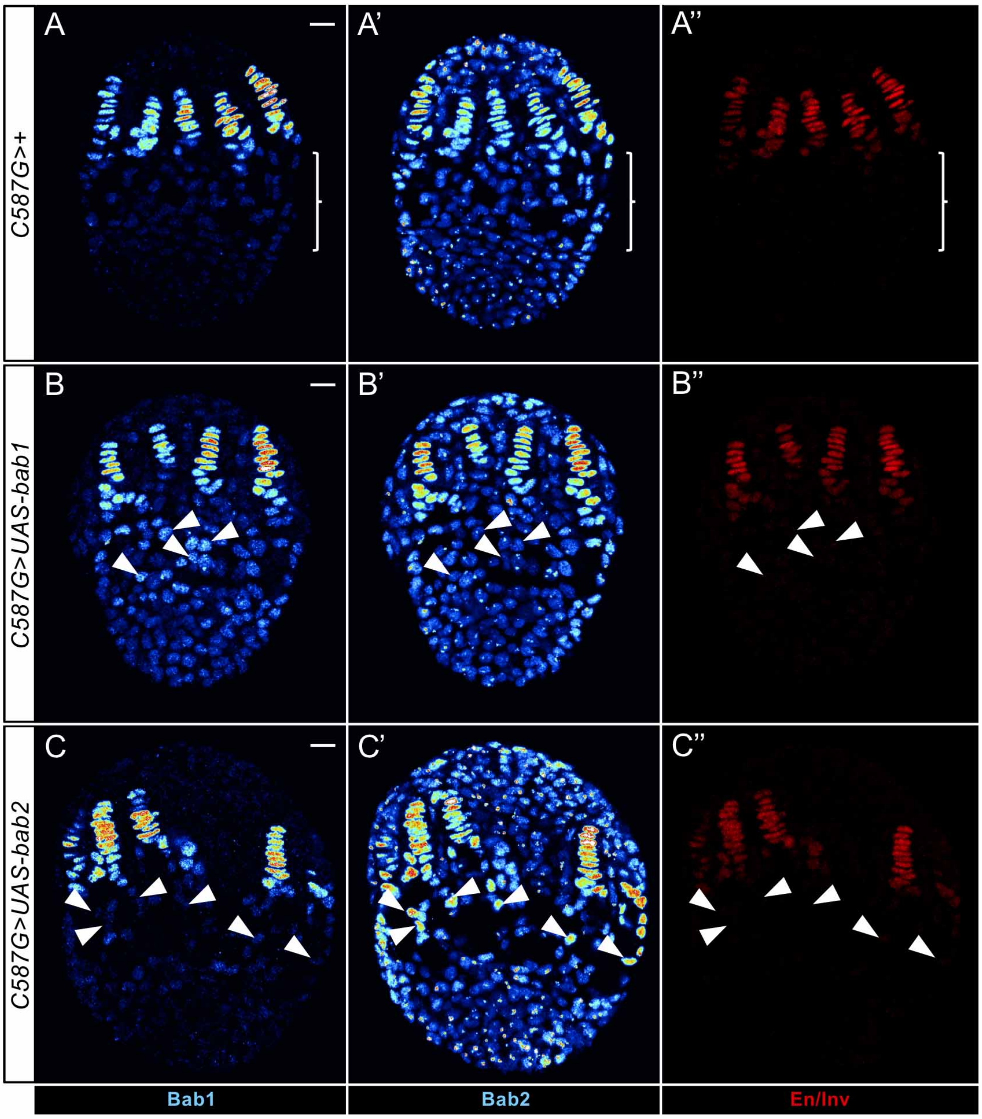
